# TandemTwister: Scalable genotyping and advanced visualization of tandem repeats

**DOI:** 10.64898/2026.01.28.702315

**Authors:** Lion Ward Al Raei, Maryam Ghareghani, M-Hossein Moeinzadeh, Martin Vingron

## Abstract

Tandem repeats are genomic regions consisting of consecutively repeated units with variable copy numbers and possible mutations. They are used in DNA fingerprinting and have been implicated in complex traits and genetic disorders, including neurodegenerative and developmental diseases. The vast and expanding number of tandem repeat loci in the human genome underscores the need for fast and scalable tools for accurate genotyping and visualization. An accurate tool for characterizing these variants is essential for understanding their functional impacts and associations with phenotypes. We developed TandemTwister, a novel algorithm implemented in C++, as a highly scalable and parallelized tool for tandem repeat copy number genotyping. Additionally, we created an interactive visualization tool to facilitate quick manual inspection, displaying exact motif occurrences, counts, and population information across haplotypes. TandemTwister demonstrates high accuracy and runtime efficiency for tandem repeat genotyping across all long-read sequencing technologies and assembled genomes. We evaluated the performance of TandemTwister in Ashkenazim trio on different sequencing technologies on a set of 1.2 million annotated tandem repeat regions. TandemTwister was the fastest and most accurate genotyping tool available for tandem repeats in comparison to the state of the art tools. For PacBio Hifi data as an example, TandemTwister was run in 17 minutes on 32 CPU cores resulting in 99.4% recall, 98.0% Mendelian consistency, and 94% sequence accuracy. We also showed a successful super-population clustering and examined inheritance patterns of tandem repeats and haplotype blocks in three trio sets. TandemTwister demonstrated its ability to detect pathogenic repeat expansions. We applied it in a cohort of 31 individuals with neurodegenerative and developmental disorders, successfully distinguishing healthy from pathogenic copy numbers.

## 1 Introduction

*Tandem repeats (TRs)* are segments in a genome made up of consecutive copies of a *repeat unit (motif)*. These repeats are categorized into two classes: *Variable Number Tandem Repeats (VNTRs)* that have a repeat unit length of at least 7 bps, and *Short Tandem Repeats (STRs)* with a repeat unit shorter than 7bps. The motif length can be as small as one single base where a single nucleotide is repeated, which is called a *homopolymer*. We consider homopolymers as a sub-class of STRs unless they are stated separately in the manuscript.

Tandem repeats are prone to mutations and to polymorphism in the repeat copy number and may have multiple alleles in the population. A TR region can consist of a single repeat unit or multiple motifs repeated interleaved with one another. To date, there is a huge (∼ 1.2 M) and still growing set of TR regions in the human genome cataloged, with size ranging from a few to several thousand base pairs [1].

*Tandem repeat genotyping* is the process of determining the alleles of a tandem repeat region by identifying the total copy number of the repeat units. For regions with multiple motifs, it additionally involves resolving the precise order and copy number of each motif within the haplotypes. Tandem repeats have been linked to various diseases such as Huntington’s disease, many ataxias, frontotemporal dementia, fragile X syndrome, amyotrophic lateral sclerosis, and other predominantly neurological disorders [2–12]. Variable number tandem repeats are thought to be a major source of contribution to “missing heritability” [13]. Characterization of tandem repeats could lead to the discovery of new associations with yet unexplained phenotypes and disease.

The functional impacts of TR variants have not been fully explored. The inherent size limitations of short-read sequencing technologies make them impractical for an accurate genotyping of long tandem repeats. The short-read-based existing methods that profile TR variations [14, 15] are limited to repeat regions that are shorter than the length of the short reads (100-200 bp) or do not provide an accurate estimation of the correct copy number for the long tandem repeat copy numbers. Long read sequencing technologies such as Pacific Biosciences (PacBio) and Oxford Nanopore (ONT) are more effective in characterizing tandem repeats because they can bridge most of the tandem repeat regions.

The current state of the art TR genotyping tools, namely *Tandem Repeat Genotyper (TRGT)* [16] and *Vamos* [17] exhibit limitations in the supported sequencing technology and accuracy in TR regions, leaving room for improvement in tandem repeat genotyping methods. There also remain opportunities to further improve computational running time, particularly when processing large datasets and massive number of regions. Additionally, the current tools lack advanced interactive visualisation for exploring and interpreting tandem repeat genotyping results.

To drive advancements in long-read-based tandem repeat genotyping, we introduce TandemTwister, a method designed for accurate and efficient tandem repeat genotyping. Comprehensive analysis on the Ashkenazi trio demonstrated TandemTwister’s superior performance in both accuracy and runtime over state-of-the-art tools. We analyzed tandem repeat variation in a population using TandemTwister on the HGSVC2 cohort and conducted trio analysis on three families within this dataset. As a proof of concept, we validated its ability to discover pathogenic repeat expansions in a cohort of 31 ONT samples from individuals with neurodegenerative and developmental disorders. In addition to the genotyping method, we developed an advanced interactive visualization tool. This tool enables manual investigation and provides deep insights into tandem repeat patterns. It includes the display options of population genotypes and pathogenic repeats. With its advanced features, the tool serves as a valuable resource for a broad target group, including biologists, geneticists, and bioinformaticians, facilitating understanding of tandem repeat patterns, structures, and variations.

## 2 Methods

*TandemTwister* genotypes tandem repeats using long DNA sequencing reads such as PacBio CLR, Hifi CCS or ONT reads, as well as genome assembly inputs. It takes as input a BAM file of the aligned long reads to the reference genome and a BED file containing the reference tandem repeat catalog including genomic coordinates with their motif sequences [1]. The output of our tool includes a VCF file containing the ordered motif occurences in each allele along with the reference genome allele.

The visualization tool is a web-based application that requires users to upload the VCF file outputted by TandemTwister, containing tandem repeat genotypes of an individual, with the optional input files for a population VCF file. It visualizes tandem repeat alleles in these regions, using color-coded motifs to represent haplotype sequences and motif occurrences.

Given the reads representing a particular TR region and its repeat motifs, *TandemTwister’s* goal is to compute the order and the copy numbers of repeat motifs for the two haplotypes. To this end, *TandemTwister* first extracts and trims reads from the annotated TR region and aligns all motifs to each of the read sequences. As a tandem repeat region can be either heterozygous or homozygous, we cluster the reads into one or two haplotype clusters according to the detected copy numbers and order of motifs. Motif alignment intervals per haplotype are then corrected and used for computing a consensus sequence of the motifs based on their occurrence in each haplotype allele.

Figure 1 shows a general overview of the algorithm workflow. It illustrates the required input data, which includes a BED file of reference TR regions with their set of motifs and a BAM file of the aligned sample sequence (either long reads or an assembled genome) to a reference genome. There is a TR border refinement step for short motifs in order to mask the reference regions that have low motif purity, then the reads are trimmed by the exact borders of tandem repeat regions (a). The illustration then represents an example of a motif alignment, which starts by computing a dynamic programming (DP) matrix of alignment of each motif to an individual cut read sequence (b). Subsequently, the best potential end-points for a motif alignment are collected (blueish rows) for each DP matrix (c). The order of motifs defined by a read sequence is then computed by a second dynamic programming algorithm that selects an optimal set of non-overlapping motif-match intervals. The next step is clustering reads into haplotypes based on their length and motif alignment features (d). It is then finalized by computing the consensus order and copy number of motifs per cluster that yields the final tandem repeat genotypes (e). The final tandem repeat genotypes are exported to a standard VCF file (**f**). This file can then be used as input for our visualization tool, which provides an advanced graphical representation of repeat regions and motif compositions (**g**). We elaborate on the different parts of the methods in the following sections.

**Fig 1.**
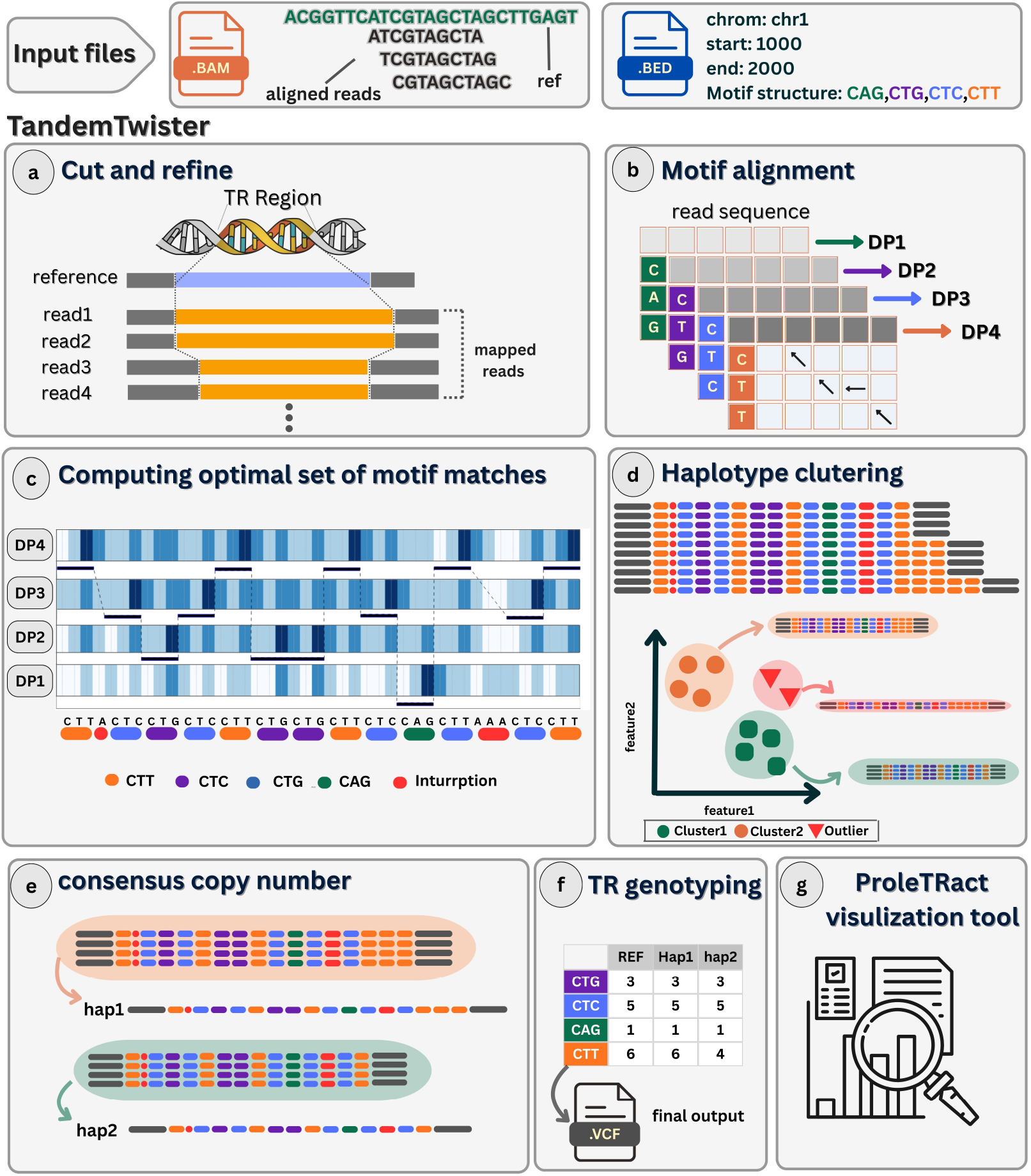
General workflow of *TandemTwister* tool. Input files: a BAM file of reads alignment to the reference genome and a BED file containing the repeats coordinates and motif structures. a) Cut and refine reads in a TR region: Colored blue and orange bars represent the cut sequence in the reference genome and reads. b) Motif alignment: Example DP matrices for aligning four motifs, represented with different colors, to a cut read. c) Each row represents a motif alignment scores for each read position. Blue color intensities represent the scores. The candidate match intervals for a read are computed from all motifs (bottom motif annotation). d) Haplotype clusteirng: example of two haplotype read groups with different motif compositions. e) Consensus TR allele computation per cluster. f) TR genotyping outputted to a final VCF file. g) ProlTRact: Tandemtwister visulization tool.

### 2.1 Preprocessing: TR border refinement

We observed many cases of annotated TRs where the motifs were distributed sparsely in the reference sequence. These cases happen because of interruption by random sequences within the repeat region. We describe such regions in terms of repeat *purity*, i.e., the fraction of the TR region covered by the annotated motifs. Although *TandemTwister* is capable of dealing with sequence interruptions, it is worth mentioning that sequencing errors in the reads, especially in regions with short motifs, make it difficult to reliably determine a consensus genotype. To address this problem, we developed a procedure to deal with motifs shorter than 5 bps by determining and then masking regions of low motif purity.

We use our motif count algorithm, which is fully described in the subsequent sections, to obtain a set of motif match intervals in every TR region in the reference genome. The next step is to compute the high-purity sub-regions of the TR region. To this end, we merge the adjacent match intervals if their distance is lower than a threshold (default: two times the maximum motif length in the region). The remaining set of merged intervals serves as the initial set of TR sub-regions. If a sub-region has a short length (less than 5bps for homopolymers and 8bps otherwise), we filter it out. Furthermore, if a remaining sub-region has a low purity (less than a threshold set to 70%), we remove the sub-region from the final set. Subsequently, the input spanning reads or assembly sequence are extracted and then trimmed in the refined TR borders. This procedure yields more precise genotyping results for all long-read technologies, especially for noisy sequencing data such as PacBio CLR and Naopore reads.

### 2.2 Motif-to-read alignment and candidate motif matches

Every TR region is annotated with a set of motif sequences. For computing the order and the number of motif matches in a read sequence, we first compute a score matrix for aligning each motif to the sequence. Afterwards, we collect all of the candidate motif match intervals based on the set of high-score positions in the sequence for all motifs. The final set with ordered motif occurrences are then computed based on a dynamic programming (DP) algorithm that we designed to find an optimal set of non-overlapping match intervals. Here we present the method details of the score matrix computation and our DP algorithm to find the best set of motif matches and their counts.

For the score matrix, we compute a dynamic programming matrix comparing the motif to a sequence. based on a Smith-Waterman-type recursion [18]. The alignment parameters are a match score of 1, mismatch penalty of 0, and gap penalty of 1. Figure 1 (middle-left) shows an example of a dynamic programming matrix for a particular pair of motif and sequence. The values in the last row of the matrix show the alignment scores ending in each of the sequence positions. We have one score matrix per motif for every read, and our goal is now to tile the sequence with motif matches derived from these dynamic programming matrices. We start backtracking from every cell in the last raw to collect candidate intervals for motif matches. Every backtracking path gives us a motif match interval that is defined as the sequence indices in the long sequence from start to the position of the maximum score in the path. The corresponding motif match interval is selected as a candidate interval if the maximum score in the backtracking path exceeds a certain threshold.

### 2.3 Computing the optimal order of motif matches

We collect the set of candidate motif match intervals for all motifs in each read. After successful completion of motifs-to-read alignment (section 2.2), each read position can be a candidate motif alignment hit where it is annotated with an aligned motif ending at that position, its match interval and the alignment score. In this section, we mention the term *interval score* several times, which refers to the alignment score of a motif match corresponding to the mentioned interval. The set of motif match intervals can be large and overlapping as illustrated in Figure 1 (part c).

We formulate the interval selection task as an optimization problem to identify the highest-scoring subset of non-overlapping intervals. More precisely, the goal is to find a subset of intervals with a maximum cumulative sum of scores with the condition that the selected intervals are not overlapping. In order to solve this problem, we have designed a dynamic programming algorithm.

We define ℐ as the set of all collected intervals and ℐ _*i*_ as the set of all intervals ending at position *i*. Each interval *I* ∈ ℐ is characterized by *S*(*I*) and *E*(*I*) as the start and end coordinates of the interval, and *W* (*I*) as its individual score. Let *n* denote the maximum end position among all intervals. For all positions *i* ∈ {1, 2, …, *n*} in the read sequence, we define ℳ (*i*) as the optimal set of non-overlapping intervals up to or before the position *i*, and *CS*(*i*) as the cumulative score of these intervals. The cumulative scores and optimal interval sets are computed using the following recursive functions:

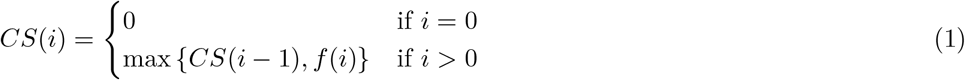

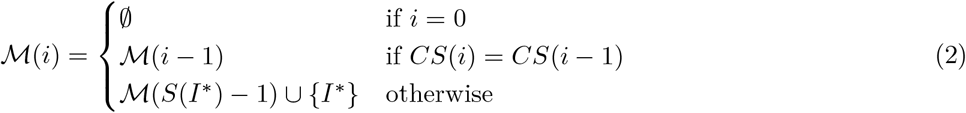

where 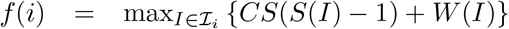 is defined as the maximum possible score over all intervals ending at position *i*, and 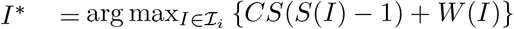 is the interval that maximizes *f* (*i*). The final solution (*n*) contains the optimal set of non-overlapping intervals ordered by their occurrence in the sequence. This algorithm is capable of finding the best set and order among multiple motifs and can also handle and detect interruptions between repeats.

### 2.4 Genotyping tandem repeats

The input of our method can be either a haploid assembled genome or a diploid sample with sequencing reads. In case of a haploid assembly input, we compute the optimal set of motif occurrences in the trimmed sequence and directly output it. For a diploid reads input, our genotyping task is to compute the TR alleles representing the order of motif occurrences in the two haplotypes, which requires clustering reads by their haplotype and computing the consensus motif orders per cluster and per tandem repeat locus. *TandemTwister* only genotypes regions that have a minimum number of spanning reads (set to 6 by default).

#### 2.4.1 Clustering reads by haplotype

In a diploid input sample, each TR region can be homozygous or heterozygous depending on the TR allele representations in the two haplotypes. Our goal is then to cluster the reads into one or two groups representing the supported haplotype alleles. To this end, we apply our motif-match algorithm for each individual read in a TR region and then cluster the reads by their haplotype based on these features: trimmed read length, the ordered set of aligned motifs, individual motif copy numbers, path purity, and intervals accumulated scores. For a refined TR region, each spanning read is separately trimmed for every selected sub-region, and the corresponding features are integrated across all sub-regions.

To remove outliers, reads whose aforementioned feature values deviated by more than 2 standard deviations from the mean were excluded from further analysis. We used DBSCAN for clustering reads by haplotype in each TR region [19], and ran it in several iterations to have a better clustering performance. DBSCAN depends on a parameter *E*, which is the radius in which a specified number of points must exist to be considered as one cluster. When a point has no neighbors within its *E* radius, it is considered noise. When we find many such noise points, we dynamically increment the *E* parameter (by 0.1) over several iterations. The iteration stops if either the number of points in the noise cluster is below a certain threshold or a maximum number of iterations (default = 40) has been reached.

We here consider a diploid human genome, and each tandem repeat region should be represented by one or two clusters, depending on whether the TR allele is homozygous or heterozygous. The result of DBSCAN, however, can be any number of clusters, therefore we further process the DBSCAN clusters. If DBSCAN gives us a cluster with at least 70% of the points, the region is considered to be homozygous with a single cluster. Otherwise, the region is heterozygous and we want to compute two clusters. To this end, we start with the two largest clusters *C*1 and *C*2 as the base clusters that represent the two haplotypes. For any potential additional cluster, each of its reads gets assigned to either *C*1 or *C*2 depending on which one it is closer to. Noise points are not considered further.

For homopolymer regions, we applied an additional criterion for heterozygosity: the two clusters must differ in length by at least 2 units, and the smaller cluster must contain a minimum fraction of reads (≥ 30% for CCS reads, ≥ 10% for ONT and CLR reads).

#### 2.4.2 Consensus tandem repeat alleles

After clustering reads by haplotype, we identify the most frequent motif-match intervals and report the read with this maximum motif match as the representative consensus read in the output VCF. For high-error reads (CLR, ONT), an optional algorithm aligns reads within a cluster to the anchor read (the previously selected representative read) and adjusts motif-match intervals. Intervals supported by at least 70% of reads are included in the consensus TR allele. The computed tandem repeat alleles are reported in the output VCF as the final genotype for each region.

## 3 Interactive visualization of tandem repeats

We developed a web-based application named *ProleTRact* for interactive visualization of tandem repeat alleles in different regions. Upon providing the input files, ProleTRact enables the analysis of tandem repeat (TR) alleles across genomic regions. It gets as input the VCF file of genotyped tandem repeats of an individual sample (generated by TandemTwister) or a directory including all corresponding sample VCFs from a population. The tool visualizes haplotype alleles with color-coded motifs, where sequence partitioning reflects motif occurrences (Figure 2). Users can navigate between adjacent regions or directly query specific TR loci using a search function. The tool has advanced interactive feature to facilitate a deep manual investigation of each tandem repeat region. The interactive features include the possibility to scroll to left and right for motif visualizations in long TR regions, displaying the repeat unit and the related metadata (e.g., sequence, copy number, GC content) by hovering over different parts of the repeat region.

**Fig 2.**
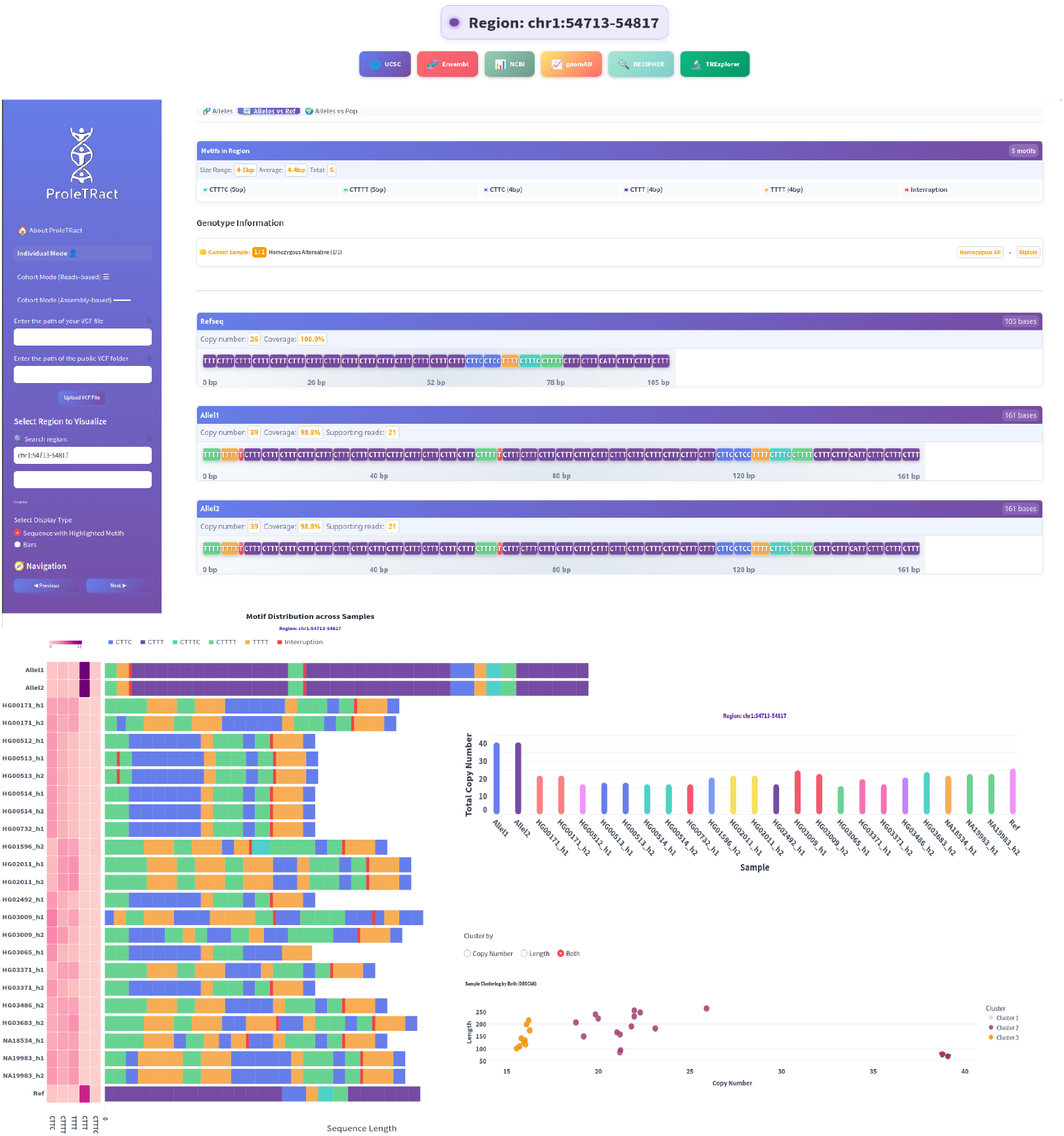
Overview of ProleTRact’s UI. The key features of the tools are displayed: (1) top-left: input options for the user to add an individual vcf file or a cohort vcf results directory and a filed for searching regions, (2) top: optional buttons for redirecting the user to differet views of the tandem repeat region in public datasets and genome browsers such as USCS, ensemble, etc. (2) top panel: visualization of tandem repeat alleles with motif-partitioned color-coding, and (3) bottom panel: population-level visualisation, including heatmaps displaying different motif counts in the population samples, stacked bar plots of motif occurences in each haplotype sample, the bar plots of total motif counts per sample and clustering samples by their motif copy number and repeat length features.

A view of the visualization application is represented in Figure 2 and supplementary figures S1-S2, displaying the application options and UI design and the tandem repeat visualizations in different modes. In order to enable fast lookups and deeper insights into public datasets and different viewers of a TR region, ProleTRact supports region lookups in public genome browsers such as UCSC, Ensemble, NCBI, genomAD, DECIPHER, and TRExplorer. By clicking on each of these websites, ProleTRact redirects the user to the specific region in the corresponding genome browser website.

ProleTRact operates in two distinct modes: 1) individual sample mode and 2) cohort mode. In individual sample mode, users provide the file path to a VCF file containing tandem repeat genotypes generated by TandemTwister for a single individual sample. Furthermore, the tool provides comparative analyses against both the reference genome and the Human Genome Structural Variation Consortium (HGSVC) population data, representing statistics on motif copy numbers and bar plots and heatmap visualizations to highlight allelic distributions. The tool also shows some geenral statistics of tandem repeats such as maximum and average motif size in the whole genome or per chromosome. In population mode, the tandem repeat genotypes of each sample are displayed along with the stacked bar plots showing the population TR alleles, and heatmaps of individual motif copy numbers across population alleles. This allows for a fast lookup into a region, observing the overall tandem repeat compositions in population, and detect potential deviations from the repeat patterns in population. There is also the option of looking at the clustered samples based on their aligned motif copy numbers and repeat length features.

For TR regions linked to disease, ProleTRact displays the pathogenic copy number threshold (see Figures 5; S6-S12), facilitating rapid assessment of whether the observed alleles fall within the healthy or disease-associated range. For annotating tandem repeat disease loci, we used STRchive [20], which is a dynamic disease-related tandem repeat catalog based on up-to-date research and clinical findings.

In summary, the visualization tool provides an interactive and comprehensive way to explore tandem repeat regions, display repeat alleles and patterns, examine population-level patterns, and identify potential deviations and pathogenic samples. Its intuitive interface makes it accessible to a broad range of users, including bioinformaticians, clinical scientists, and geneticists.

## 4 Results

We present a comprehensive analysis of the results of *TandemTwister* in comparison with the other state of the art tools in different sequencing technologies in the Ashkenazi trio (HG002, HG003, HG004). We further present our results in the population analysis of a cohort of 35 genomes from Human Genome Structural Variation Consortium (HGSVC2) [21] and demonstrate the population clustering and trio haplotype blocks based on tandem repeat profiles. As a clinical applications, we elaborate on our proof of concept for detecting pathogenic repeat expansions in a cohort of Nanopore disease samples with neurodegenerative and developmental disorders [22].

### 4.1 Comparison with other tools

We ran *TandemTwister* (v0.1.0) on a set of reference tandem repeat annotations (≈ 1.2 million regions [23]) on the Ashkenazi trio samples HG002 (child), HG003 (father), and HG004 (mother) on different sequencing technologies PacBio CLR, HiFi CCS, and Nanopore reads. We compared our genotyping results with *TRGT* (v4.0.0) and *Vamos* (v2.1.7), which are the state of the art tools that have analogous input and output. The input to the tools is an alignment BAM file and a reference set of tandem repeat region coordinates with their motif sequence, and the output includes the genotyped tandem repeats in these regions, that is the order of motif occurrences in each allele.

The first difference between the tools is their support for different long-read technologies. The *TRGT* tool is designed solely for PacBio HiFi sequencing reads, and does not support the noisy long read inputs such as PacBio CLR and Nanopore reads. However, *TandemTwister* and *Vamos* support all of these technologies.

Figure 3 represents an overview of a comparative analysis between the tools over various measures including running time, Mendelian consistency, and sequence concordance with high quality haplotype assembly.

**Fig 3.**
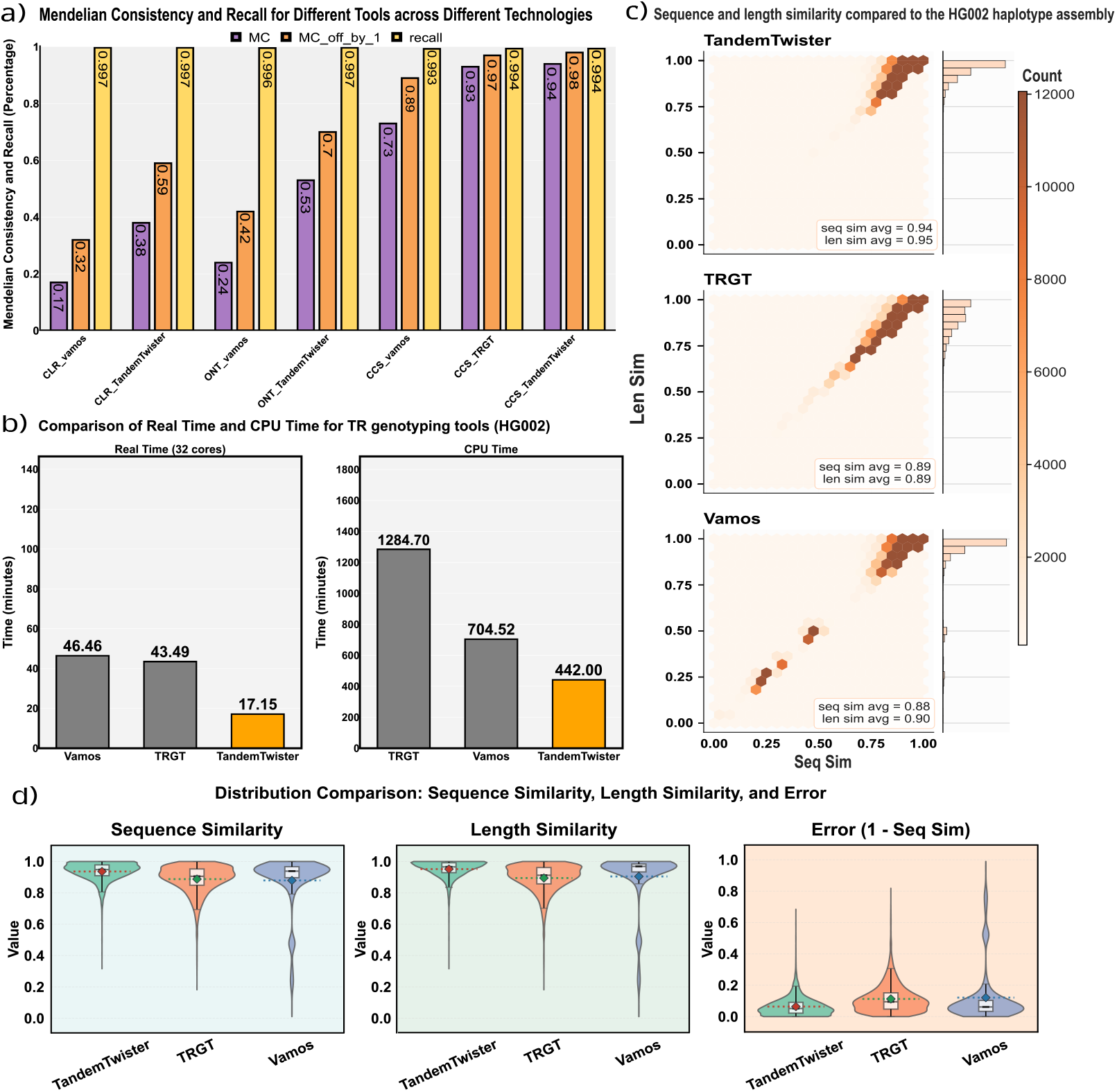
Evaluation of TR genotyping tools. a) Mendelian consistency (MC) and recall values for different long-read sequencing technologies and TR genotyping tools. Purple and Orange bars show the default and off-by-one Mendelian consistencies in the Ashkenazi trio, respectively, and yellow bars represent the recall values (fraction of the genotyped regions) for the child sample HG002. b) bar plot illustration of running time performance comparison of the tools based on their wall-clock time (using 32 cores) and CPU running time. The fastest genotyping tool is *TandemTwister*. c) Comparison of the concatenated TR allele sequences with the high-quality haplotype assemblies. The analysis is for the CCS sample HG002 and different subplots show performances of various tools. Each subplot shows the length and sequence similarity distribution across all TR regions per tool and sequencing technology. d) Comparison of different tool’s distributions of sequence similarity (left), length similarity (middle) and sequence error rate (right).

#### 4.1.1 Recall and Mendelian consistency

Mendelian consistency in the context of tandem-repeat genotyping is an essential evaluation metric for ensuring the reliability and accuracy of genetic analyses. The evaluation of tools based on Mendelian consistency allows for the identification of potential errors and biases.

We call a tandem repeat region *Mendelian consistent* if the child inherits the exact motif copy number alleles from both parents, so following this definition, the measured *Mendelian consistency (MC)* is the fraction of tandem repeat regions that are Mendelian consistent. In order to use a secondary and more lenient measure, an *off-by-one Mendelian consistent* region is defined for allowing at most one copy number deviation between the child’s TR motif copy numbers and the inherited parental alleles. The *off-by-one Mendelian consistency* measure is hence defined accordingly to be the proportion of such regions in all tandem repeat loci.

Figure 3a shows the comprehensive results on comparison of the Mendelian consistency of the three tools on all sequencing technologies. *TandemTwister* genotyped 99.4% of the TR regions on CCS samples with a Mendelian consistency of 94% and off-by-one MC of 98.0%. Recall and Mendelian consistency (default and off-by-one) for CCS reads were 99.4%, 93.0%, and 97.0% in *TRGT* and 99.3%, 73.3%, and 89% in *Vamos. TandemTwister* has slightly higher Mendelian consistency measures over *TRGT* (improved MC by 1%) and is clearly better than *Vamos* (21% and 9% improved MC and MC-off-by-one).

We also evaluated the performance of both *TandemTwister* and *Vamos* on PacBio CLR and Nanopore reads and compared their performance. The improvement over Vamos was also observed on CLR and ONT reads (between 21% to 28% improvements in all four measures of Mendelian consistency). It should be noted that *TRGT* tool is specifically designed to process CCS reads, and hence, it was only used for comparison with CCS reads.

#### 4.1.2 Running time analysis

We compared the real wall-clock time and CPU time performance of *TandemTwister, TRGT*, and *Vamos* on CCS reads of the HG002 sample (Figure 3b). *TandemTwister* was the fastest tool with 442 CPU minutes, which is 1.6× faster than *Vamos* and 2.9× faster than *TRGT*. The wall-clock time using 32 cores was 17.15 minutes for *TandemTwister*, which was at least 2.5 × faster than both *Vamos* and *TRGT*. The running times analysis suggests that *TandemTwister* is the fastest TR genotyping tool for long-read sequencing, which makes it the most scalable tool for large-scale data and population tandem repeat genotyping.

#### 4.1.3 Sequence accuracy

Sequence quality and accuracy is an important and essential measure to explore the performance of the tools. In order to assess the accuracy of the genotyped sequences in TR regions, we first reconstructed the TR sequence alleles by concatenating the reported set of constituting motifs and then compared the resulting sequence and length similarity of the reconstructed alleles to the high quality haplotye assemblies of the diploid Genome in the Bottle (GIAB) sample HG002 [21]. Sequence and length similarity together reflect the concordance of the computed repeat alleles with the ground truth assembly sequences. We observed that *TandemTwister* has the highest performing measures of sequence and length accuracy with average values of 0.94 and 0.95, respectively, showing a compelling improvement of 5% to 6% over both *Vamos* and *TRGT* (Figure 3c,d). An analogous comparison has been performed between *TandemTwister* and *Vamos* on CLR and ONT reads in which we observed similar results. Our genotyping tool has improvement on sequence and length similarity in both CLR and ONT reads compared to *Vamos* (see Supplementary Figures S3-S4).

In summary, comprehensive analysis on Ashkenazim trio with different measures such as Mendelian consistency, running time and sequence accuracy suggest that *TandemTwister* outperforms the other tools.

### 4.2 Analysis of tandem repeats in population

To examine the applicability of tandem repeat profiles for population clustering and determining haplotype inheritance patterns, we analysed TR copy numbers in a population of assembled haplotype samples from the HGSVC2 cohort [21]. The cohort consists of 35 samples with 70 high-quality assembled haplotypes. The samples come from 6 different super populations: African (AFR), Admixed American (AMR), East Asian (EAS), European (EUR), and South Asian (SAS). For this analysis, we used a subset of 816, 927 tandem repeat regions that were genotyped in all samples by TandemTwister. The cohort also contains three family trios which we analysed to show how TR variations identify haplotype inheritance patterns.

Figure 4a shows the two-dimensional PCA plot of the tandem repeat copy numbers. The points are individual sample haplotypes that are colored by their super population. The results show an expected clustering of samples according to their super-populations. The African population (AFR), positioned on the right of the plot, is clearly separated from the others and exhibits higher variance along *PC*1. On the left, the East Asian (EAS) population forms a distinct and moderately dispersed cluster. The remaining three super-populations are located closer together, with South Asian (SAS) samples slightly separated and European (EUR) and Admixed American (AMR) samples co-clustered. This clustering pattern is consistent with established population structure inferred from global human genomics diversity studies, such as the 1000 Genomes Project [24] and the haplotype ancestry block analysis in the HGSVC2 study [21], both of which report substantial European haplotype ancestry within the AMR super-population.

**Fig 4.**
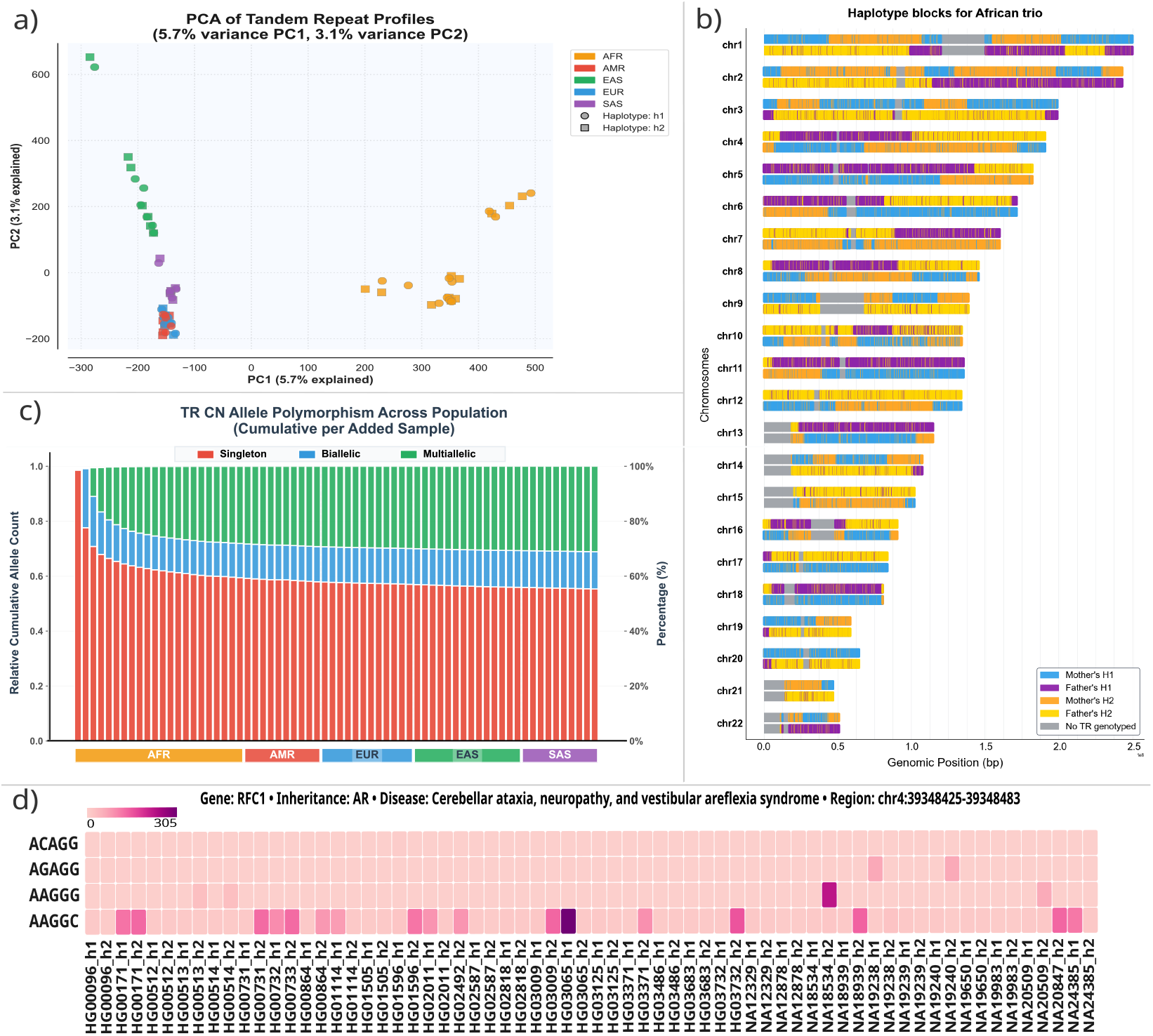
Tandem repeats in population. a) A two-dimensional PCA plot of the tandem repeat copy numbers in HGSVC2 assembly samples. samples are colored by their super population. b) Inherited haplotype alleles from the parents in African trio. Different colors refer to the mother or father alleles that are inherited to the child haplotype h1 or h2 (four possible combinations). c) The TR regions are divided into three categories by their copy number variation: Singleton (red), biallelic (blue), and multi-allelic (green). The bar plot shows the cumulative portion of these categories of tandem repeats in population by adding samples one by one in the displayed order of super-populations. d) heatmap of motif counts in hgsvc2 population in the RFC1 gene composed of four different motifs.

We further investigated the haplotype inheritance patterns in the three HGSVC2 trio families with African, East Asian and Admixed American ancestries. Figure 4b shows the results of haplotype inheritance analysis for the African trio (child: NA19240, mother: NA19238, father: NA19239). Since the full chromosome-wide haplotype assemblies were available, we were able to assign the child’s H1 and H2 haplotypes to the corresponding parent for each chromosome. It is worth mentioning that the assembled haplotypes are pseudo-haplotypes meaning that the true parent is not determined, whereas the parental haplotypes are fully determined and distinguished per chromosome. The homozygous regions of parents and the set of *de novo* tandem repeats are excluded and not shown in the haplotype inheritance analysis. The analysis shows an expected pattern of few numbers of relatively large haplotype blocks within each chromosome and parent. These results support only a few crossovers per chromosome, which is consistent with the low likelihood of the crossover events. We made an analogous analysis of the inherited haplotype blocks for the other trios, the results of which are shown in Supplementary Figure S1. We observed consistent haplotype block inheritance patterns in the other two trios supporting only a few crossover events per chromosome.

In order to study tandem repeat polymorphisms in population, we examined the number of singleton, biallelic and multi-allelic tandem repeats by their copy number profiles by adding the population haplotype samples one by one in order of their super population. As evident in Figure 4, there is a sharp drop of singletons and rise of biallelic and multi-allelic variants at the start of the sample-adding experiment. The change in the proportions of these variants continues to consistently move towards more biallelic and multi-allelic tandem repeats with a smoother alteration rate. The resulting analysis does not show a saturation point, suggesting the possibility of detecting more variations and polymorphisms in population by increasing the sample size. Including all samples, 55% of tandem repeat regions are singleton and have the same copy number in the whole population, leaving a significant portion of loci with 14% biallelic and 31% multi-allelic variants.

We performed a similar analysis of tandem repeats based on their other characteristics: length and sequence. The observed patterns were analogous to those seen for copy number, with the main difference being a lower number of singletons (49% for length feature and 45% for sequence feature) and a higher number of biallelic and multi-allelic variations (Figures S5-S6, Supplementary Material). These differences are consistent with our expectation that different ways of characterizing tandem repeats capture varying levels of resolution and population diversity. For example, defining tandem repeats by their exact sequence provides the finest resolution, enabling more refined variant discovery and a higher number of distinct alleles. For TR sequence alleles, we used one-hot encoding to generate numeric features, under the assumption that all sequence alleles of a given TR region should be treated as equally distant from each other..

As an example of a variable tandem repeat region, Figure 4d shows the motif copy numbers in the RFC1 gene across the population. This gene contains four highly similar motifs, and TandemTwister accurately computes the copy numbers for each motif (Section 4.3). Abnormal expansions on specific motifs in RFC1 are associated with cerebellar ataxia, a neurological disorder affecting balance and movement functions. The corresponding heatmap illustrates the variation in motif copy numbers in this region across the HGSVC2 population, highlighting its natural benign variation among healthy individuals.

### 4.3 Pathogenic tandem repeats

In order to validate the ability of *TandemTwister* to detect pathogenic repeat copy numbers, we tested our tool on a cohort of 31 targeted Nanopore sequencing samples with neurological pathogenic STR expansions. The data has been published in a recent study [22], from which we chose the set of experimentally validated genes *FMR1, HTT, ATXN1, DAB1, FXN, PABPN1, DMPK*, and *NOTCH2NLC* with their pathogenic STR expansions. *TandemTwister* successfully genotyped all 31 samples for all of the studied genes. It was done by accurately identifying STR expansions that met or exceeded the pathogenic thresholds. These thresholds were predetermined for each of the studied genes based on existing literature [20, 22].

Figure 5 shows the result of *TandemTwister* in the repeat expansion of *RFC1* gene, responsible for Cerebellar ataxia, neuropathy and vestibular areflexia syndrome (CANVAS). CANVAS is an autosomal recessive neurodegenerative disease, which arises from a pathogenic repeat expansion. It is specifically caused by an excess of 500 or more of “AAGGG” repeat within the *RFC1* gene. This condition is fully penetrant when motif count of “AAAGG” reaches 40 or more. Expanded STR alleles in RFC1 are relatively common and exist in several different motif conformations. The motifs “AAAAG” and “AAAGG” are considered nonpathogenic, regardless of their counts. In addition to the canonical pathogenic motif “AAGGG”, a rare “ACAGG” motif and mixed “AAAGG–AAGGG” conformation are both considered pathogenic ([25]), while the pathogenicity of various other observed conformations is currently unknown. Figure 5 displays the reported copy numbers by *TandemTwister* for AAGGG,ACAGG,AGGGC,AAGGC,AGAGG repeats across 31 samples. For example, for the disease sample R210023, *TandemTwister* reports a total copy number of 1335 and 1100 for the two haplotypes that exceeds the pathogenic threshold. *TandemTwister* was also successfully applied to detect the pathogenic repeat expansions in the other genes, the results of which are presented in Figures S6-S13 in the supplementary materials. All of these visualization plots for pathogenic samples are generated by ProleTRact, our interactive visualization tool for tandem repeats.

**Fig 5.**
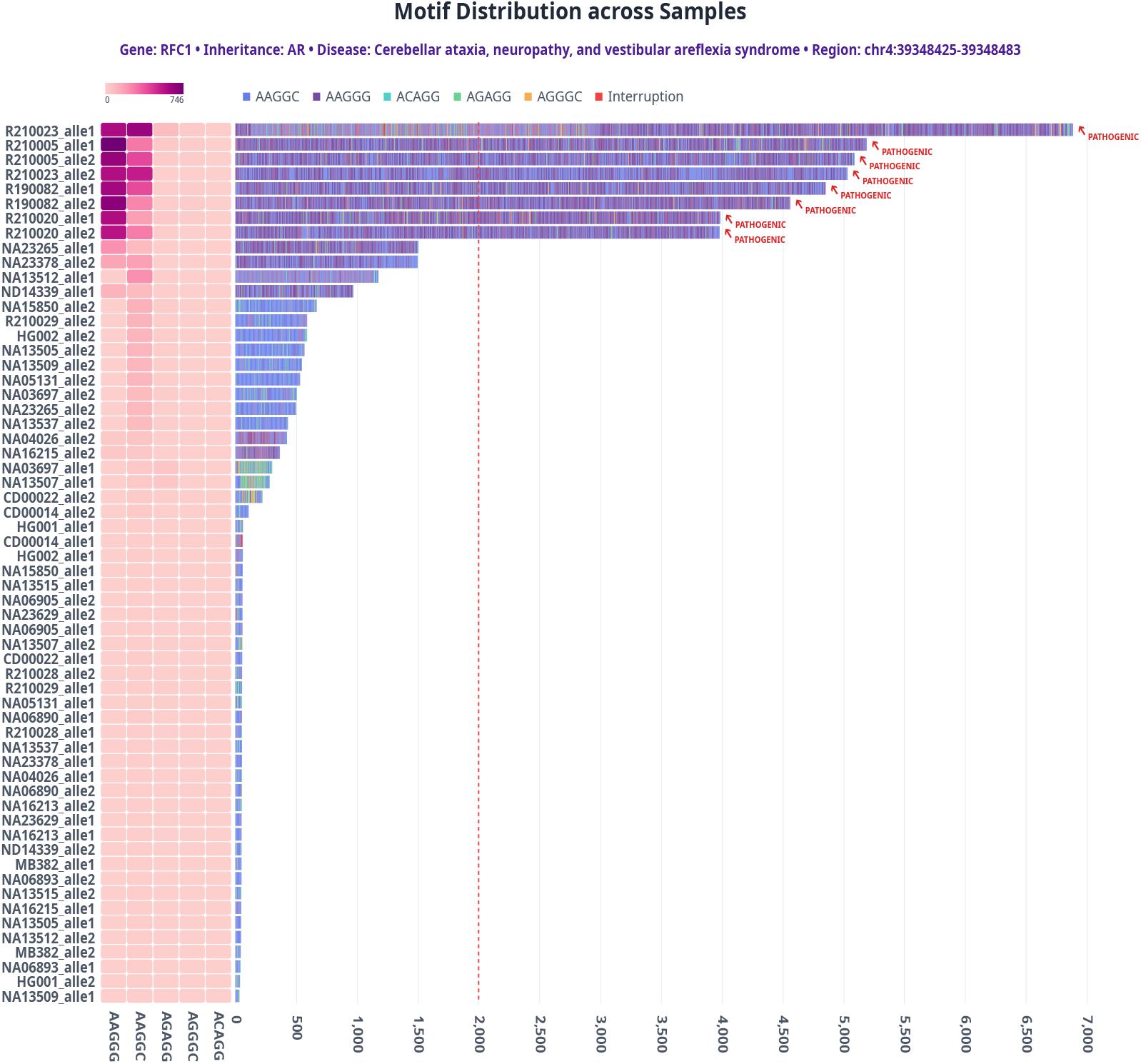
Combined visualization of *TandemTwister* genotyping results for the RFC1 gene across 31 samples. The left panel shows a heatmap displaying motif counts per sample, and the right panel shows an stacked bar plot of motif occurrences. Each sample has two bars representing its two copy number alleles. The pathogenic copy number cutoff is shown by a red vertical dashed line in the bar plot.Samples with alleles higher than the pathogenic cutoff are considered affected (4 samples; 8 alleles in total).

## 5 Discussion

We presented *TandemTwister*, a highly scalable and fast tool for genotyping tandem repeats. It produces high quality results not only for accurate long reads (PacBio HiFi CCS) but also for noisy sequencing reads of PacBio CLR and Nanopore ONT technologies. *TandemTwister* can also support a phased assembled genome as input. The tool is based on several methodological innovations and novel algorithms with a series of computational steps including TR region purification, motif alignment, motif counting, haplotype clustering, computing the repeat alleles, and making the final consensus genotypes. It is implemented as a fast, efficient and parallelized C++ code, and works with the standard generic input/output formats, e.g., bam, bed, and vcf files.

We compared *TandemTwister* with the other state of the art TR genotyping tools and showed superiority of our tool in all measured evaluation metrics including sequence accuracy, Mendelian consistency and running time. The determined tandem repeat profiles are highly informative with respect to population studies, which we illustrated by analysing TR copy numbers in a population cohort of 35 samples with 70 assembled haplotypes [21]. A clustering of the samples by TR copy number reproduced an expected super population clustering. We additionally investigated the three trio families and reconstructed the inherited haplotype blocks based on their TR copy number alleles. We further validated our tool on a set of experimentally-validated pathogenic STR expansions in a cohort of 31 ONT samples. All of the pathogenic tandem repeat expansions were successfully detected and confirmed in the corresponding samples using our genotyping tool.

In addition to our fast and accurate genotyping tool, we also proposed an advanced interactive visualization tool that enables exploring tandem repeat regions at differnt views. It has individual and cohort modes with various interactive plots and visualisations inclusing stacked bar plots of ordered motif occurrences, repeat count heatmaps and reported TR statistics. It also shows the existing genotypes of samples and the pathogenic repeat expansion thresholds based on a dynamic catalog of disease-related tandem repeats [20]. The tool enables visual investigation in individual tandem repeat regions, which facilitates the path for new discoveries for tandem repeat patterns and detecting new potential abnormalities and disease-causing variants. The tool is a web-based application with an intuitive UI design making it practical and easy to use for a wide range of researchers including bioinformaticians, experimental scientists and clinicians.

The high scalability and speed of *TandemTwister* make it suited for many biological applications, such as genotyping large sets of tandem repeats in big population cohorts and patient samples. It enables further deep investigation of tandem repeats, their population structure, and their contribution to various diseases and phenotypes. Exploration of tandem repeats can go beyond the human genomes and include different organisms, which opens a direction of research into the evolution of these variants in multiple species.

## Code and data access

*TandemTwister* is written in C++ and is available at https://github.com/Lion-ward/TandemTwister. The source code and instructions for our visualisation tool, *ProleTRact*, is also publicly available at https://github.com/Lionward/ProleTRact. We used the tandem repeat catalog from the public dataset [23]. The PacBio CLR, HiFi CCS, and Nanopore sequencing data for the Ashkenazi trio were obtained from 13 the GIAB GitHub repository [26]. The assembly data utilized in the population analysis are available on the HGSVC website [21]. The samples for the pathogenic tandem repeat expansions are derived from NCBI under the accession number PRJNA786382.

## Supplementary material

All supplementary materials, including figures, are provided in a single document.

## Acknowledgments

We would like to express our appreciation to Nico Alavai and Jakob Hertzberg for useful scientific discussions and the IT team at Max Planck Institute for Molecular Genetics for their support with technical aspects related to this project.

## Author contributions

M.G., H.M., and M.V. conceived and conceptualized the research problem. M.G. and H.M. performed initial experiments during the early phase of the project. Building on this initial work, M.G. and L.W.A.R. designed the current version of the algorithm. L.W.A.R. implemented the TandemTwister C++ code and the visualization tool. M.G. and L.W.A.R. designed and carried out the data analysis experiments. The manuscript was primarily written by M.G., with contributions from L.W.A.R. Figures and graphical representations were prepared by M.G. and L.W.A.R. The manuscript was edited and polished by M.V. All authors read and approved the final manuscript.

## Declaration of competing interests

H.M. holds the position of CTO at Lucid Genomics and M.V. serves as an advisor for Lucid Genomics. These roles have been disclosed to ensure transparency in the reporting of this research.

## Notes

### Summary of Updates

To revise the name of one of the authors.

