## Supplementary figures for "TandemTwister: Scalable genotyping and advanced visualization of tandem repeats"

### Supplementary Information for: [TandemTwister: Scalable genotyping and advanced visualization of tandem repeats]

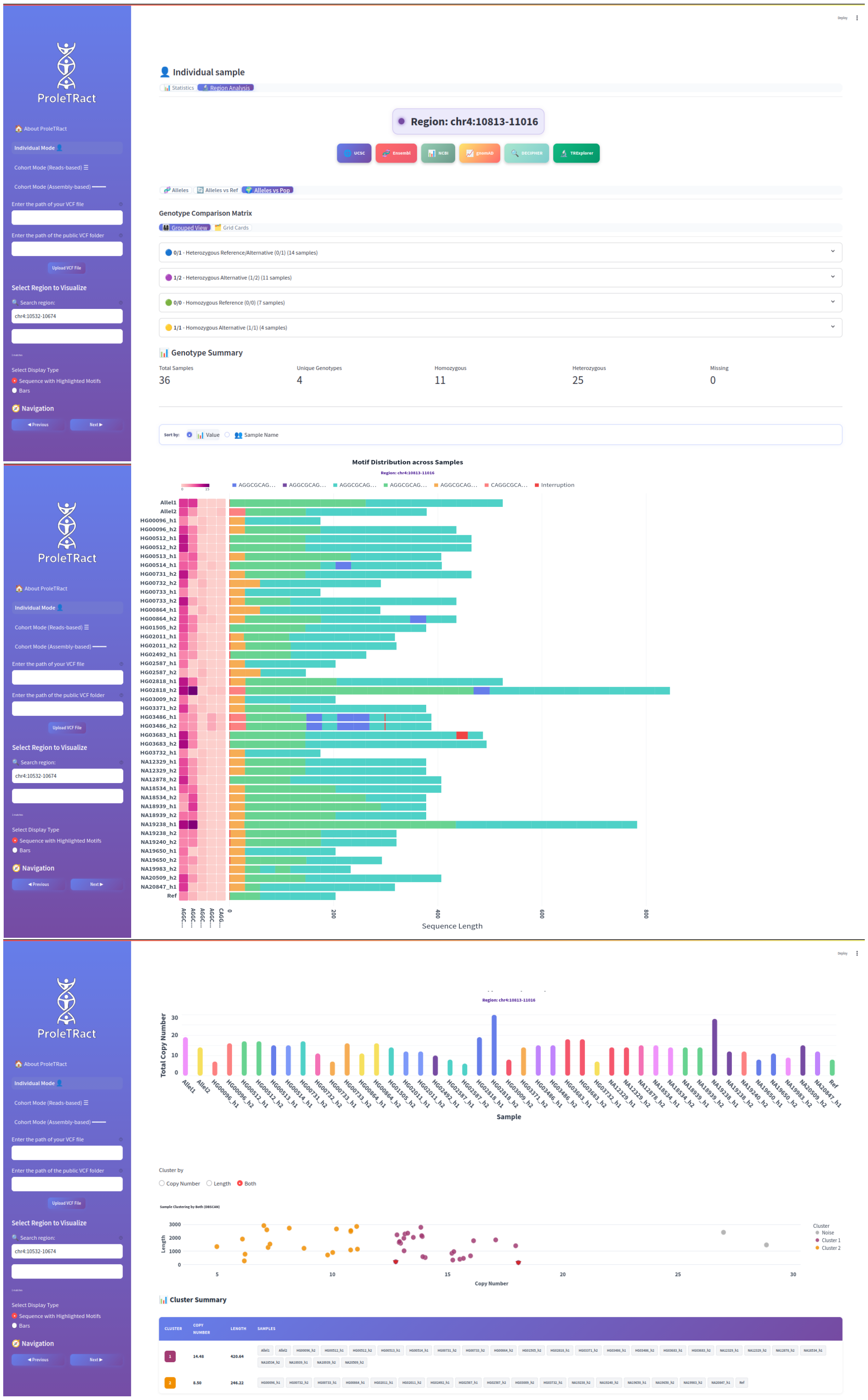


Figure S1: Overview of ProleTRact's UI. The following features are displayed: Top panel: optional buttons for redirecting the user to differet views of the tandem repeat region in public datasets and genome browsers such as USCS, ensemble, etc. Information and groups of genotypes for popualtion samples, together with summary counts for uniques, homozygous and heterozygous samples. Middle panel: population-level visualisation, including heatmaps displaying different motif counts in the population samples, stacked bar plots of motif occurences in each haplotype sample. Bottom panel: population-level visualisation, including heatmaps bar plots of total motif counts per sample and clustering samples by their motif copy number and repeat length features. A clustering summary is also provided by listing the names of samples falling into different clusters and their corresponding copy number and length range.

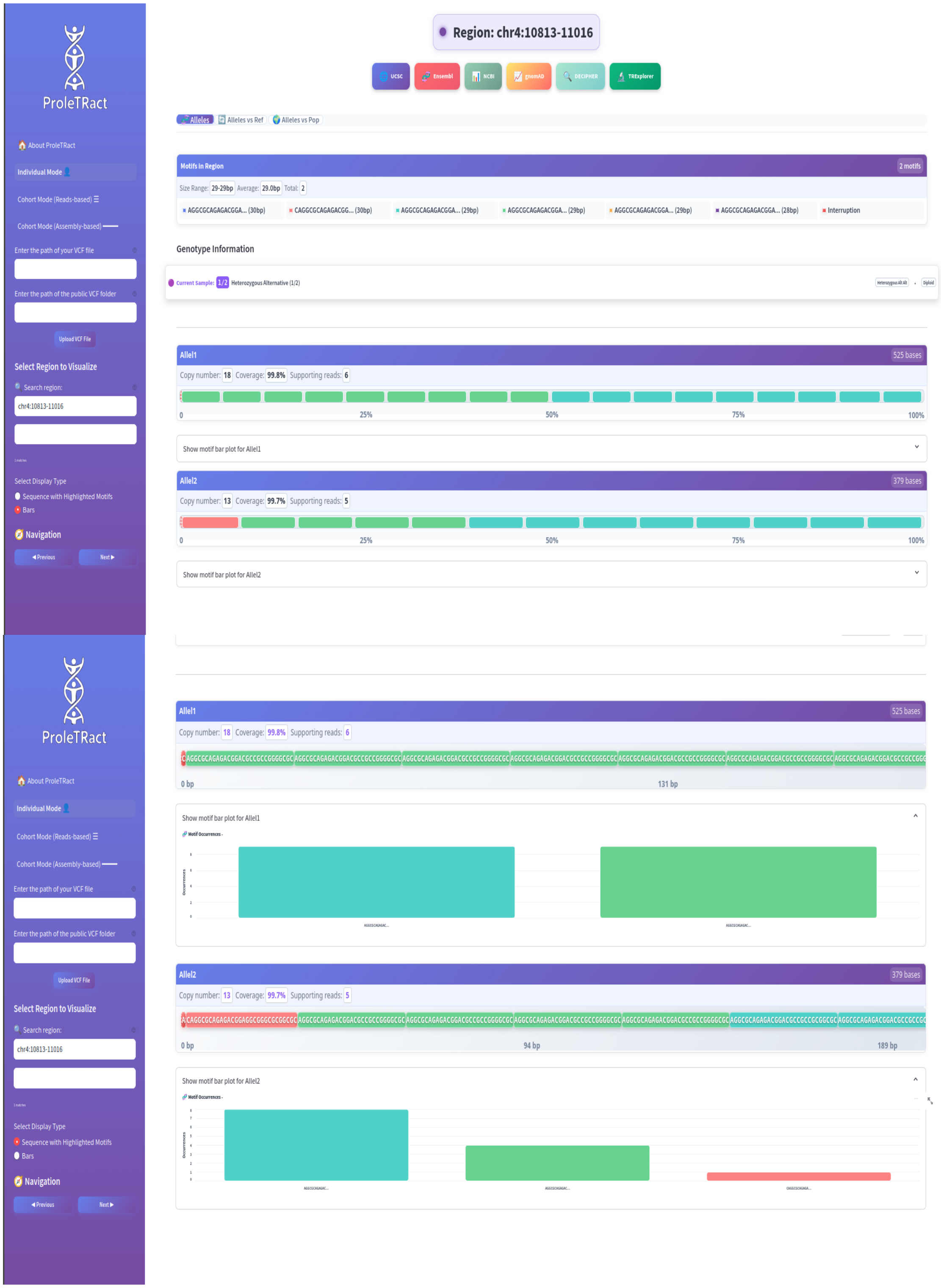

Figure S2: Overview of ProleTRact's UI. The following features are displayed: Top panel: optional buttons for redirecting the user to differet views of the tandem repeat region in public datasets and genome browsers such as USCS, ensemble, etc. Visualisation of motif compositions in tandem repeat alleles by colored bars including the totl copy number and read coverage information. Bottom panel: Color-coded bar plots representing individual motif copy numbers of each allele.

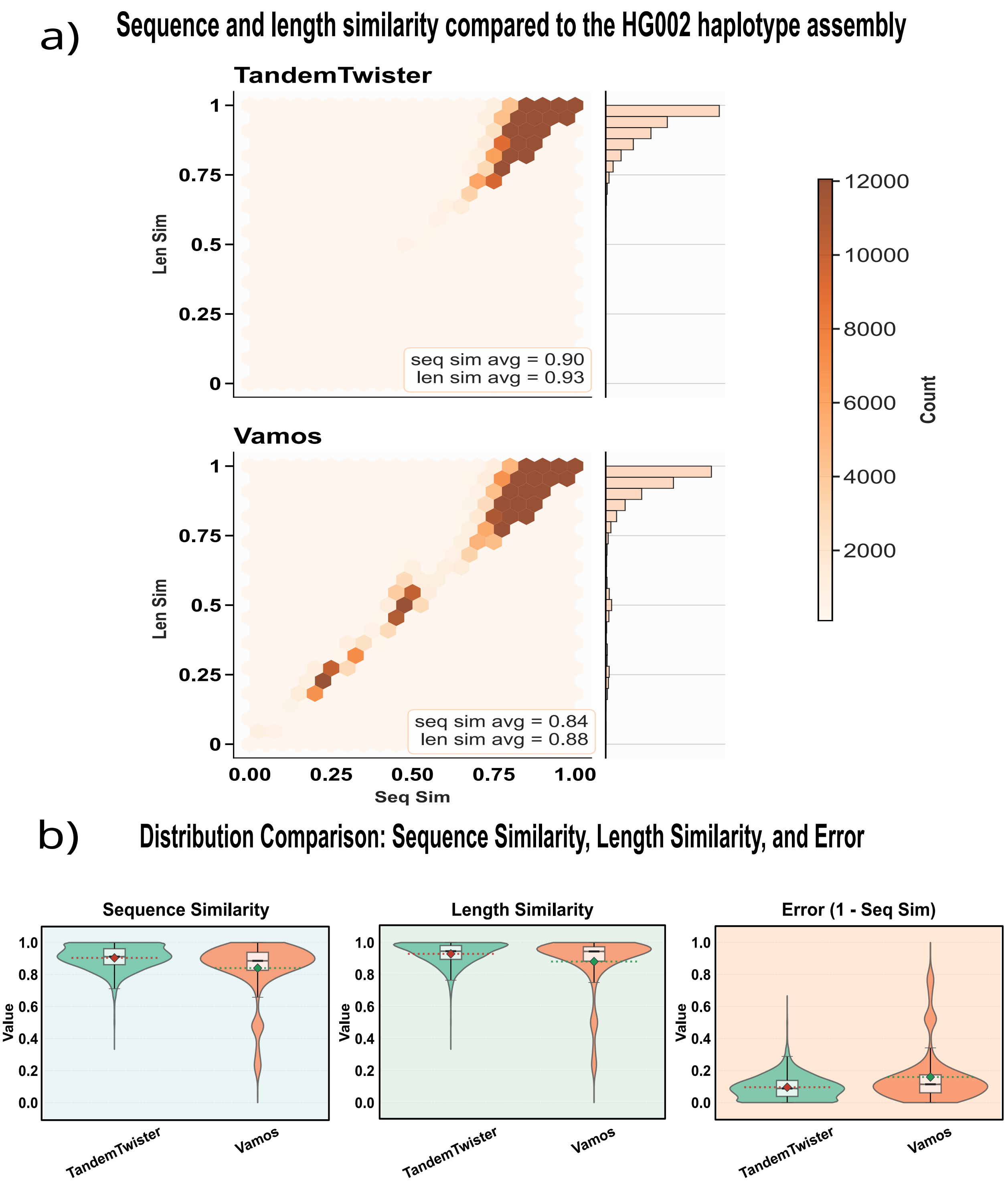


Figure S3: Sequence accuracy on PacBio CLR sample HG002. a) Comparison of the concatenated TR allele sequences with the high-quality haplotype assemblies. Hexa plots representing density of existing regions with various sequence and length similarities in the two tools. d) Comparison of different tool's distributions of sequence similarity (left), length similarity (middle) and sequence error rate (right).

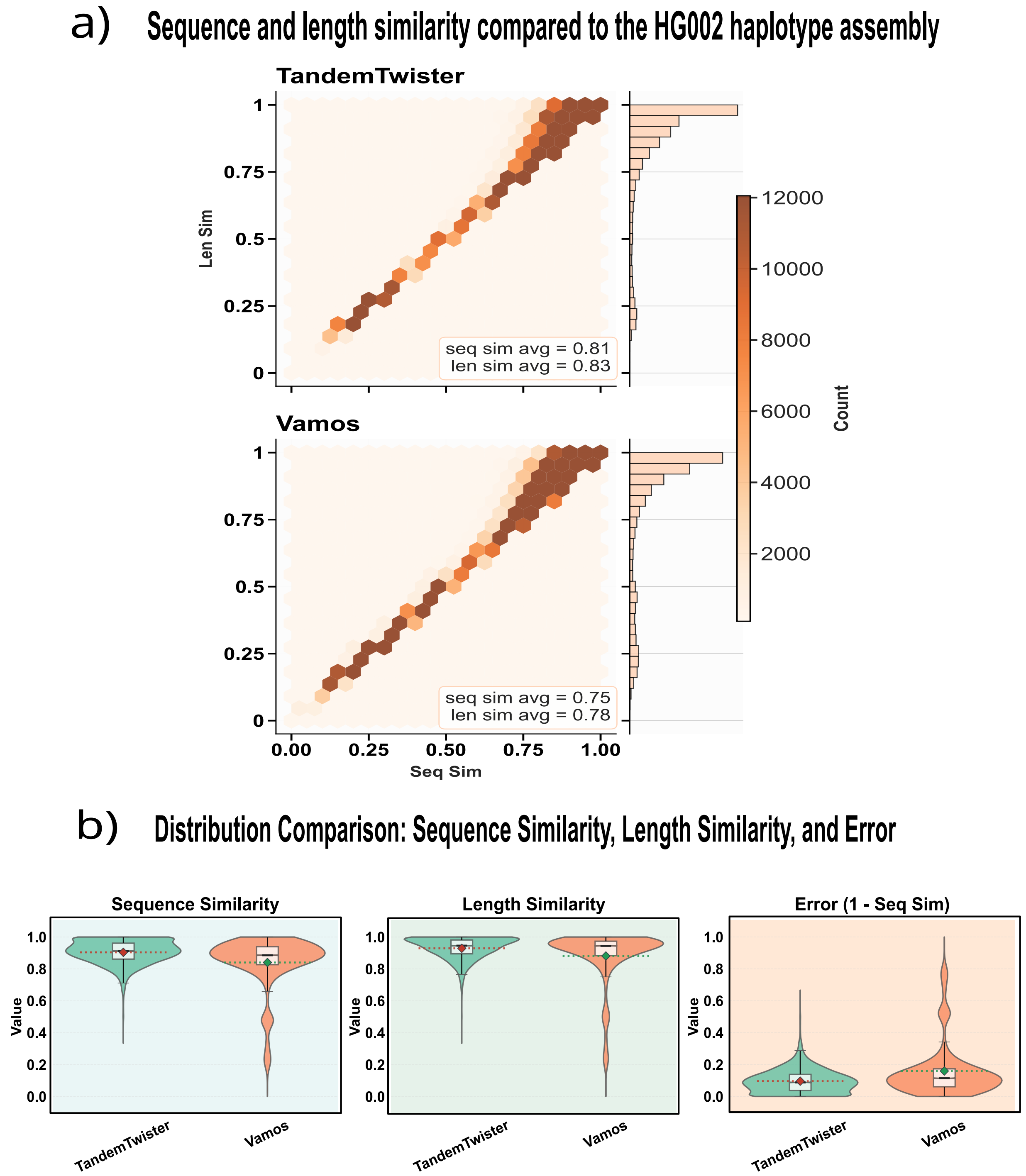


Figure S4: Sequence accuracy on Oxford Nanopore (ONT) sample HG002. a) Comparison of the concatenated TR allele sequences with the high-quality haplotype assemblies. Hexa plots representing density of existing regions with various sequence and length similarities in the two tools. d) Comparison of different tool's distributions of sequence similarity (left), length similarity (middle) and sequence error rate (right).

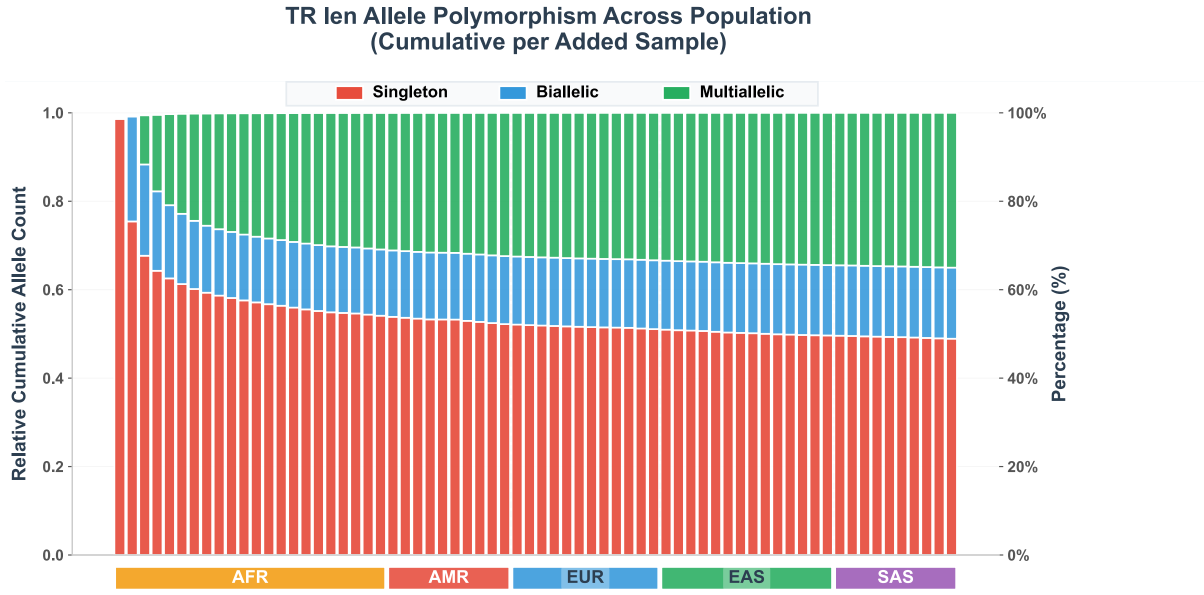


Figure S5: Accumulative plots on tandem repeat variation by length in population. The TR regions are divided into three categories by their length variation: Singleton (red), biallelic (blue), and multi-allelic (green). The bar plot shows the cumulative portion of these categories of tandem repeats in population by adding samples one by one in the displayed order of super-populations.

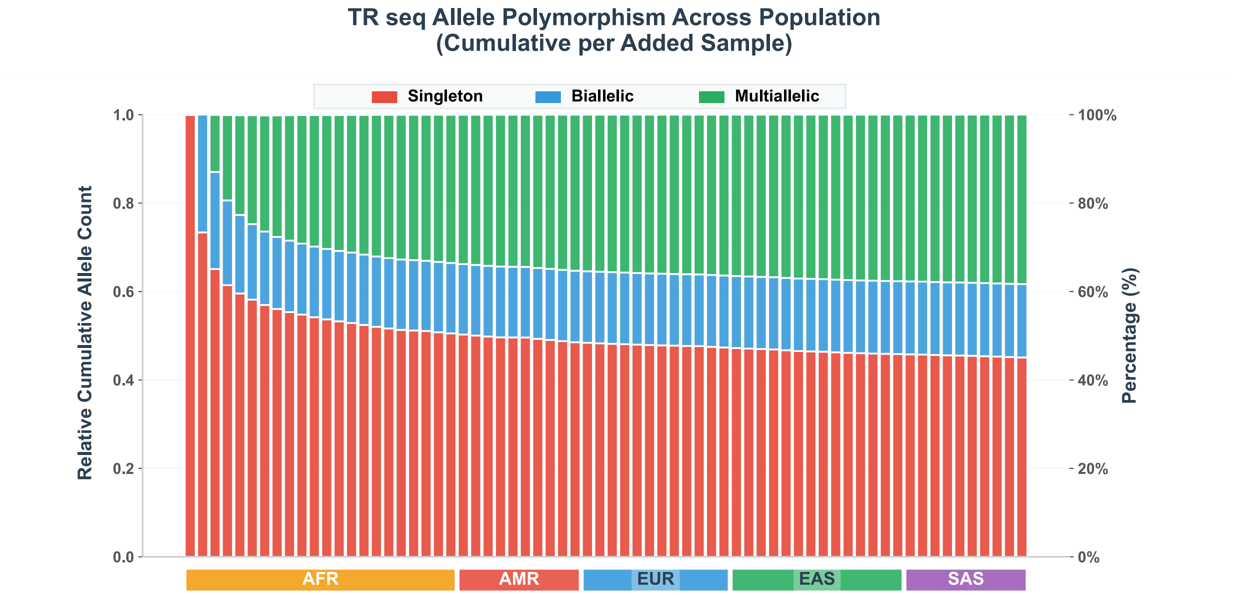


Figure S6: Accumulative plots on tandem repeat variation by sequence in population. The TR regions are divided into three categories by their sequence variation: Singleton (red), biallelic (blue), and multi-allelic (green). The bar plot shows the cumulative portion of these categories of tandem repeats in population by adding samples one by one in the displayed order of super-populations.

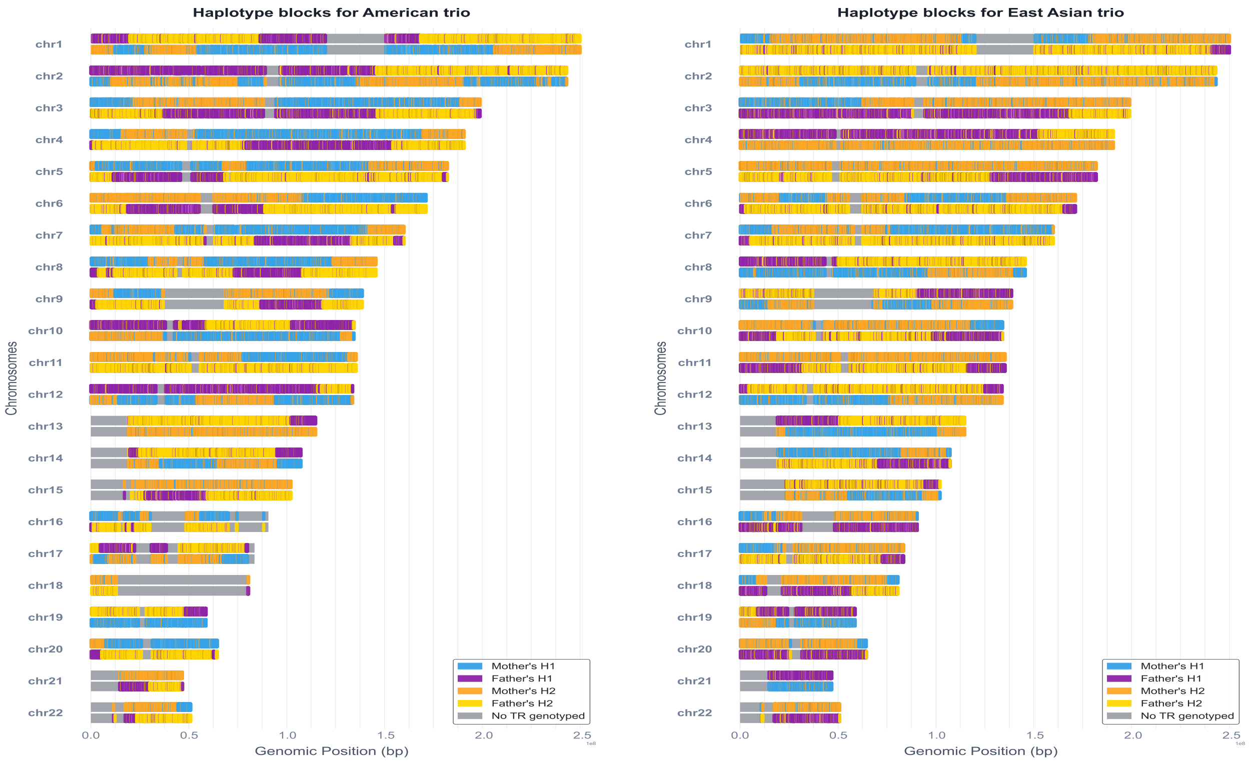


(a) Mixed American trio inheritance pattern

(b) East Asian trio inheritance pattern

Figure S7: Inherited haplotype alleles from parents in different population trios. a) Admixed American (AMR) trio, b) Ease Asian (EAS) trio. Different colors refer to the mother or father alleles that are inherited to the child haplotype h1 or h2 (four possible combinations)

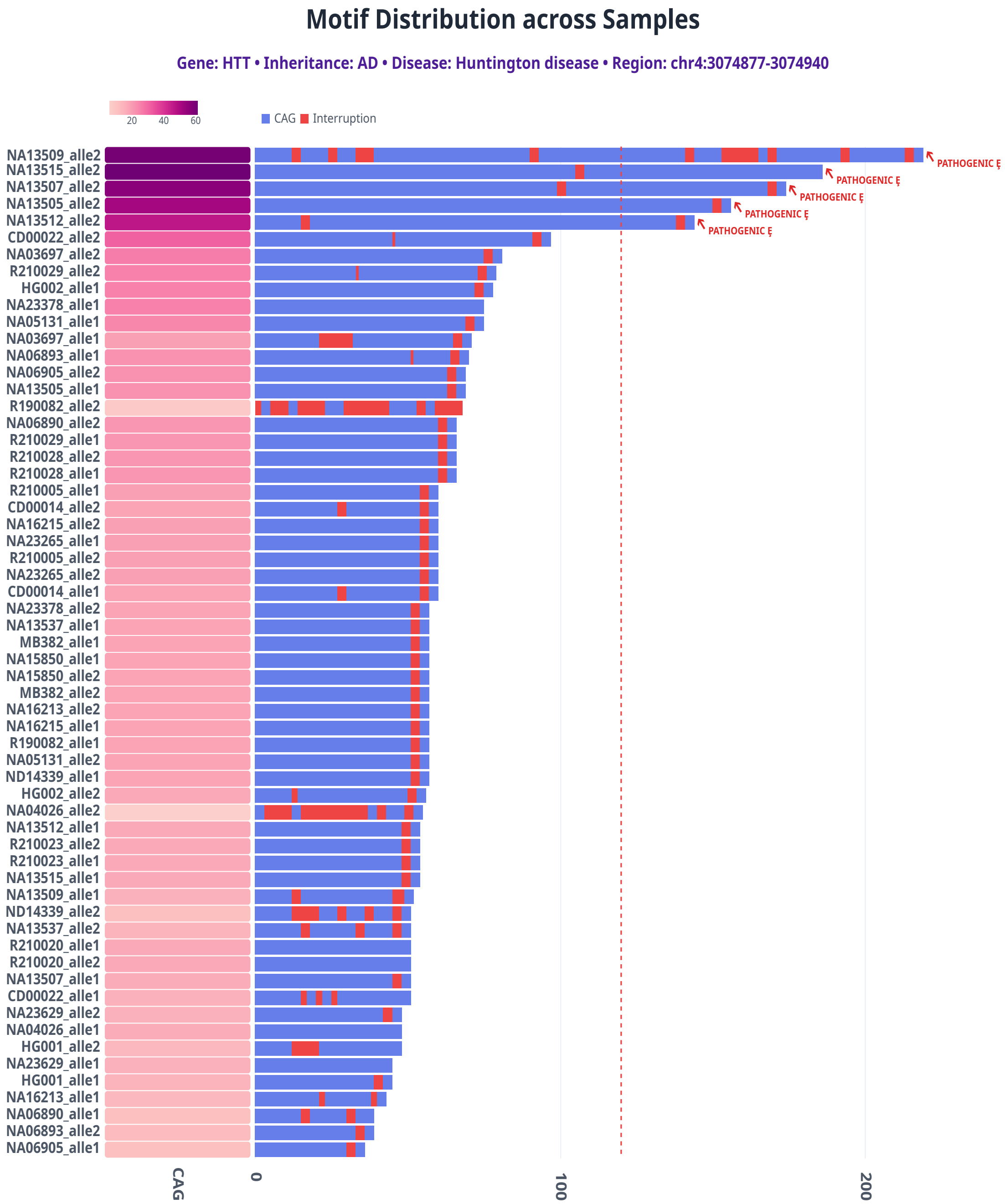


Figure S8: TandemTwister genotyping results for HTT gene across 31 samples. Left: Heatmap of motif counts; Right: Bar plot showing copy number alleles per sample. Red dashed line indicates pathogenic cutoff. Five samples are affected by the disease and show pathogenic expansion.

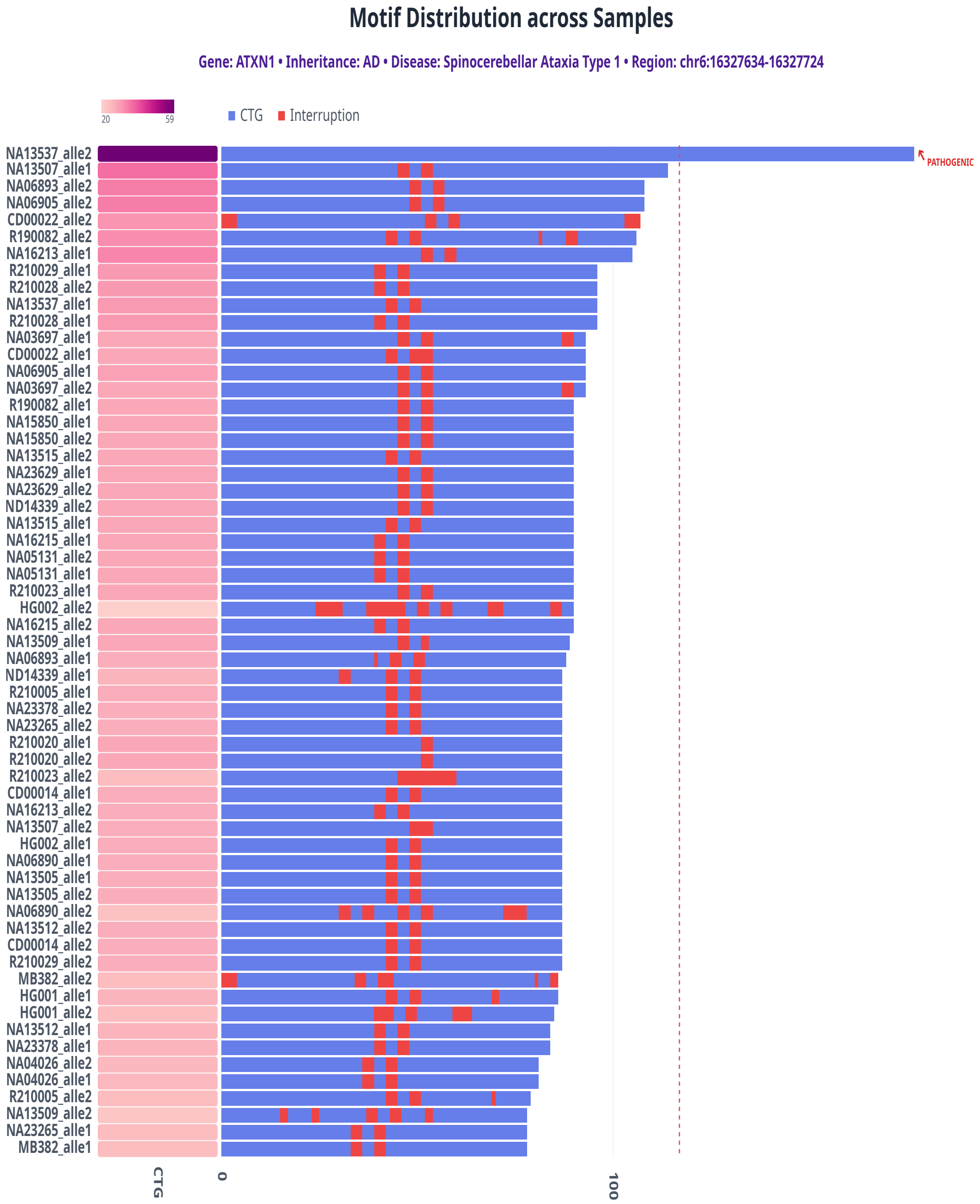


Figure S9: TandemTwister genotyping results for ATXN1 gene across 31 samples. Left: Heatmap of motif counts; Right: Bar plot showing copy number alleles per sample. Red dashed line indicates pathogenic cutoff. Sample NA13537 shows pathogenic expansion.

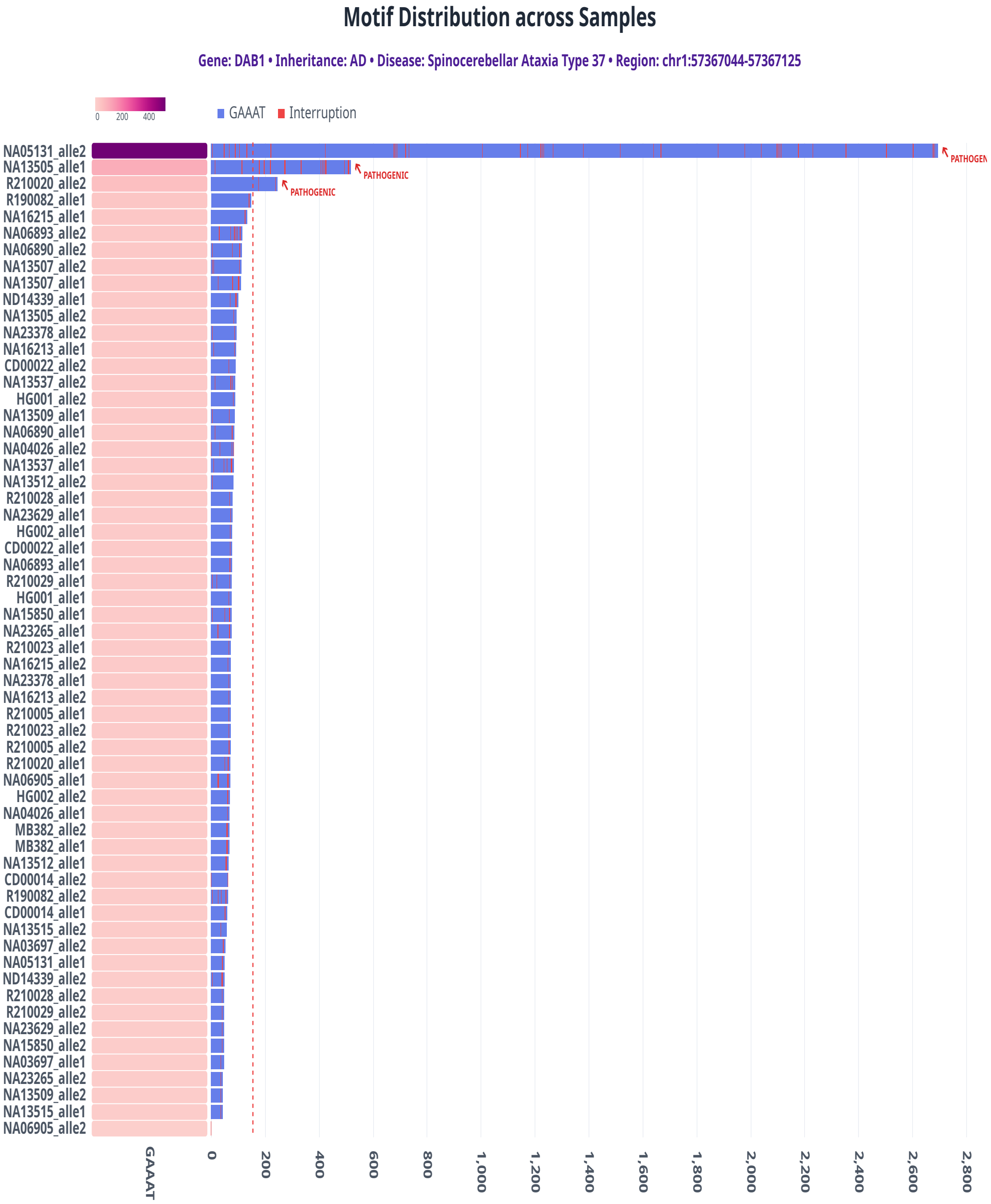


Figure S10: TandemTwister genotyping results for DAB1 gene across 31 samples. Left: Heatmap of motif counts; Right: Bar plot showing copy number alleles per sample. Red dashed line indicates pathogenic cutoff. Three samples are affected by the disease and show pathogenic expansion.

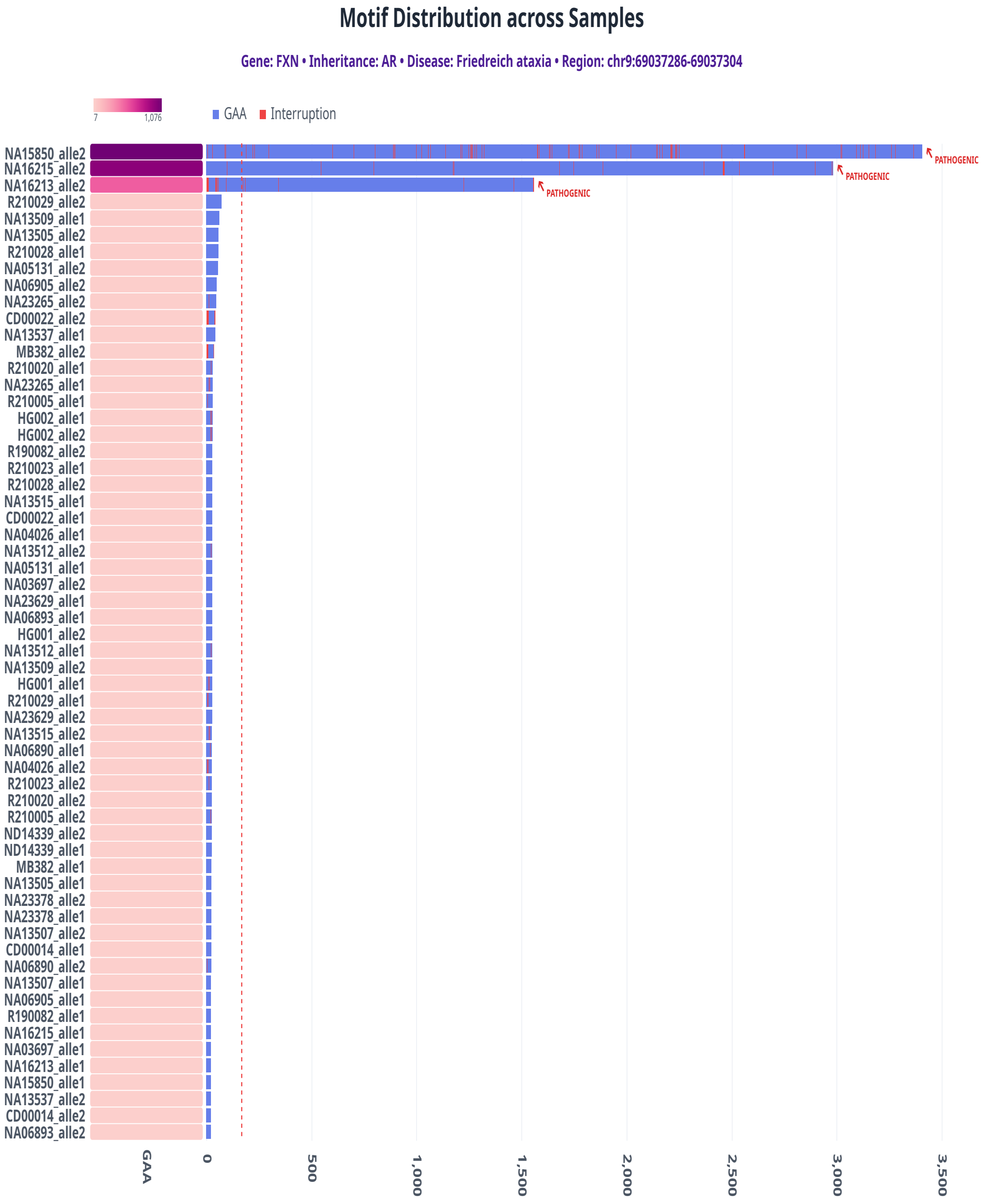


Figure S11: TandemTwister genotyping results for FXN gene across 31 samples. Left: Heatmap of motif counts; Right: Bar plot showing copy number alleles per sample. Red dashed line indicates pathogenic cutoff. Three samples exhibit pathogenic expansions above the cutoff.

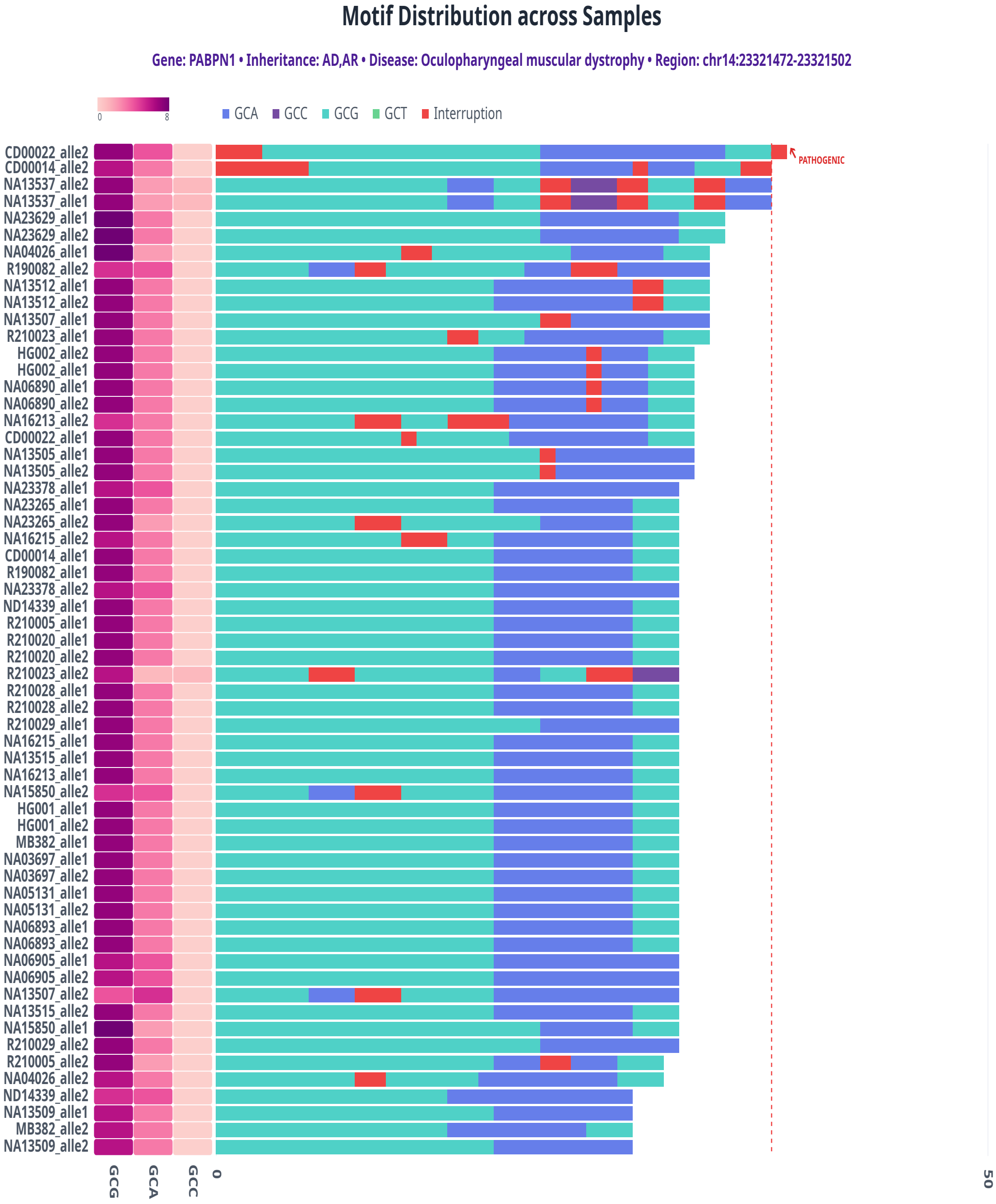


Figure S12: TandemTwister genotyping results for PABPN1 gene across 31 samples. Left: Heatmap of motif counts; Right: Bar plot showing copy number alleles per sample. Red dashed line indicates pathogenic cutoff. Sample CD00022 shows pathogenic expansion.

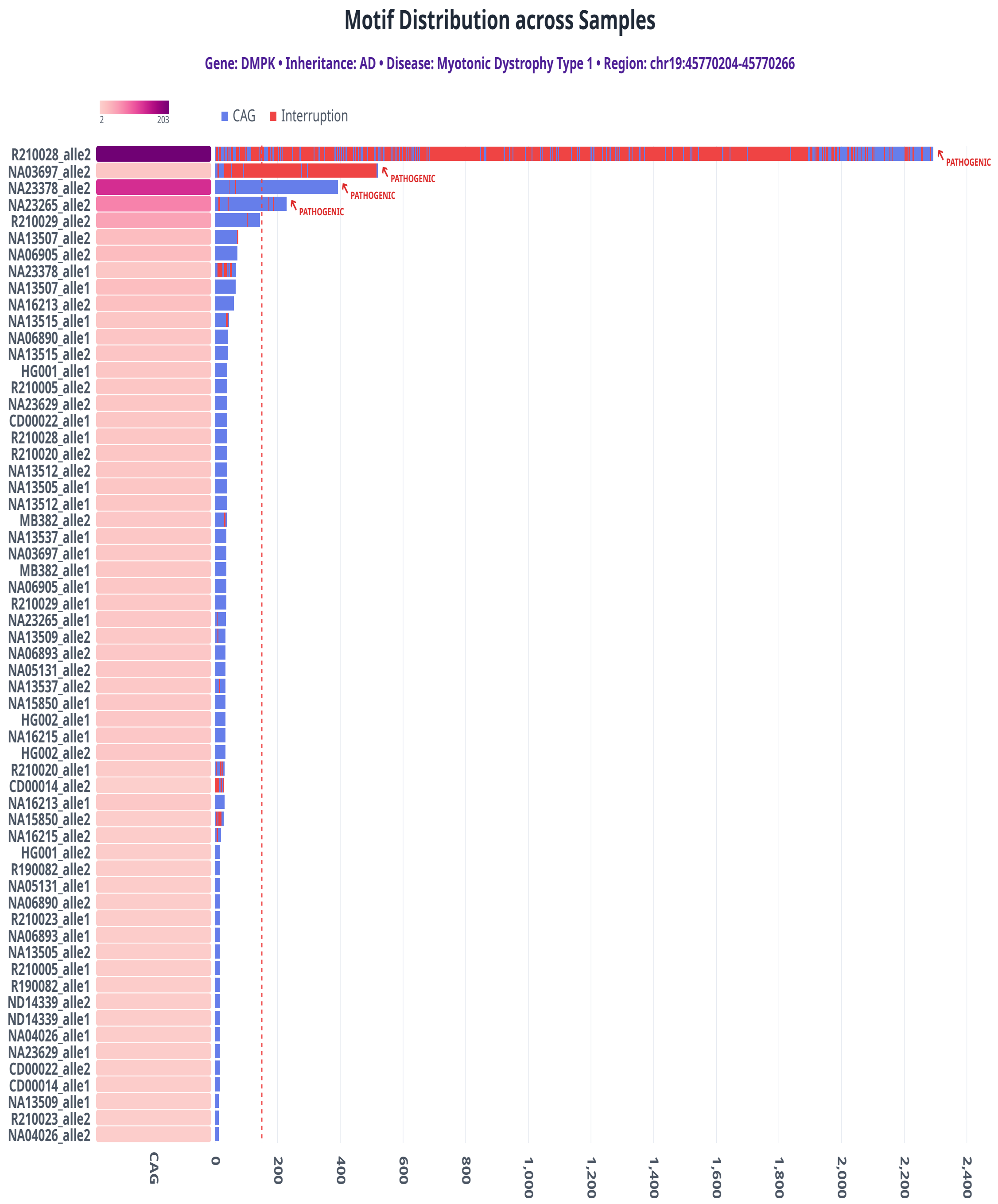


Figure S13: TandemTwister genotyping results for DMPK gene across 31 samples. Left: Heatmap of motif counts; Right: Bar plot showing copy number alleles per sample. Red dashed line indicates pathogenic cutoff. Four samples are affected by the disease and show pathogenic expansion.

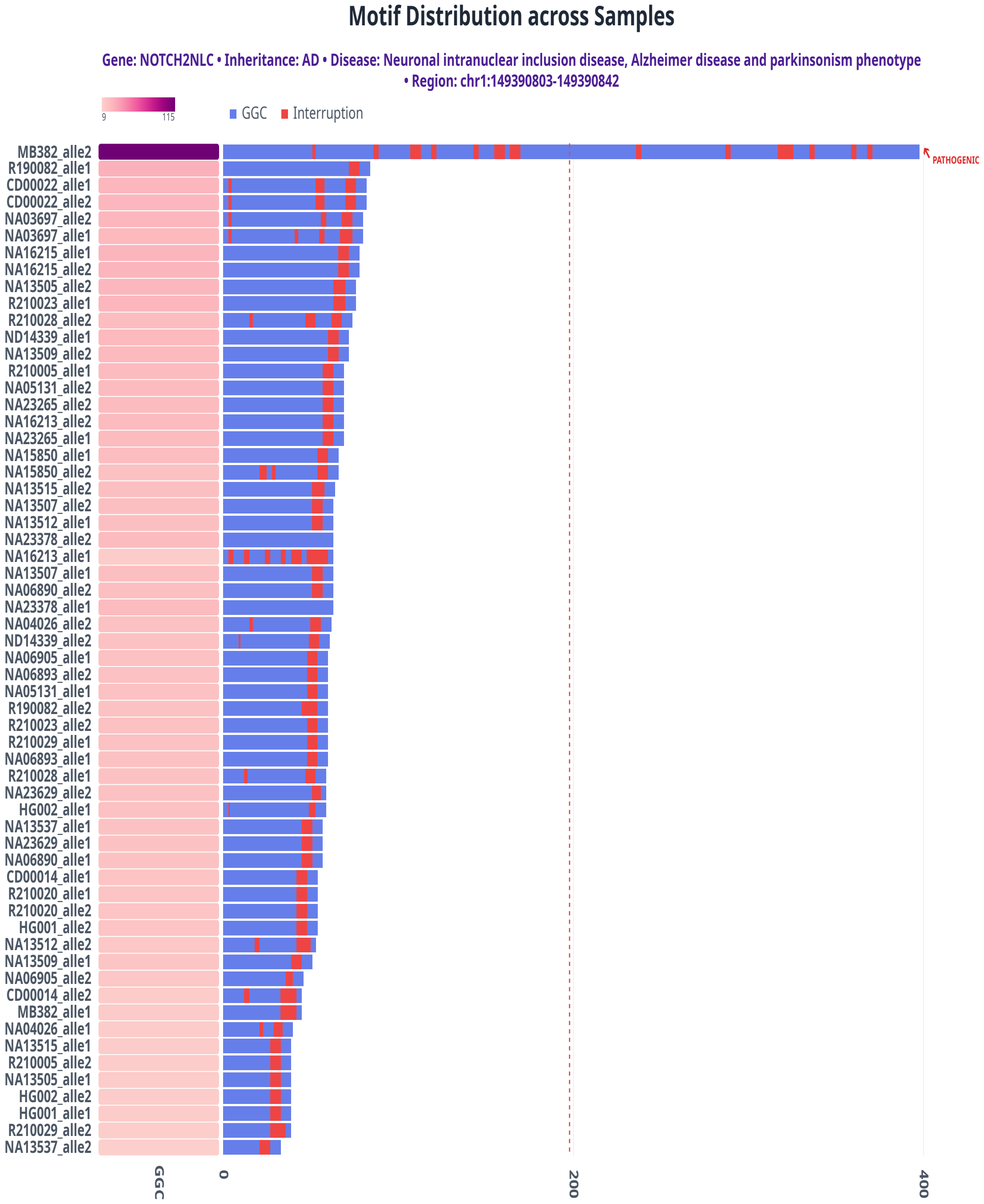


Figure S14: TandemTwister genotyping results for NOTCH2NLC gene across 31 samples. Left: Heatmap of motif counts; Right: Bar plot showing copy number alleles per sample. Red dashed line indicates pathogenic cutoff. Sample MB382 shows pathogenic expansion.

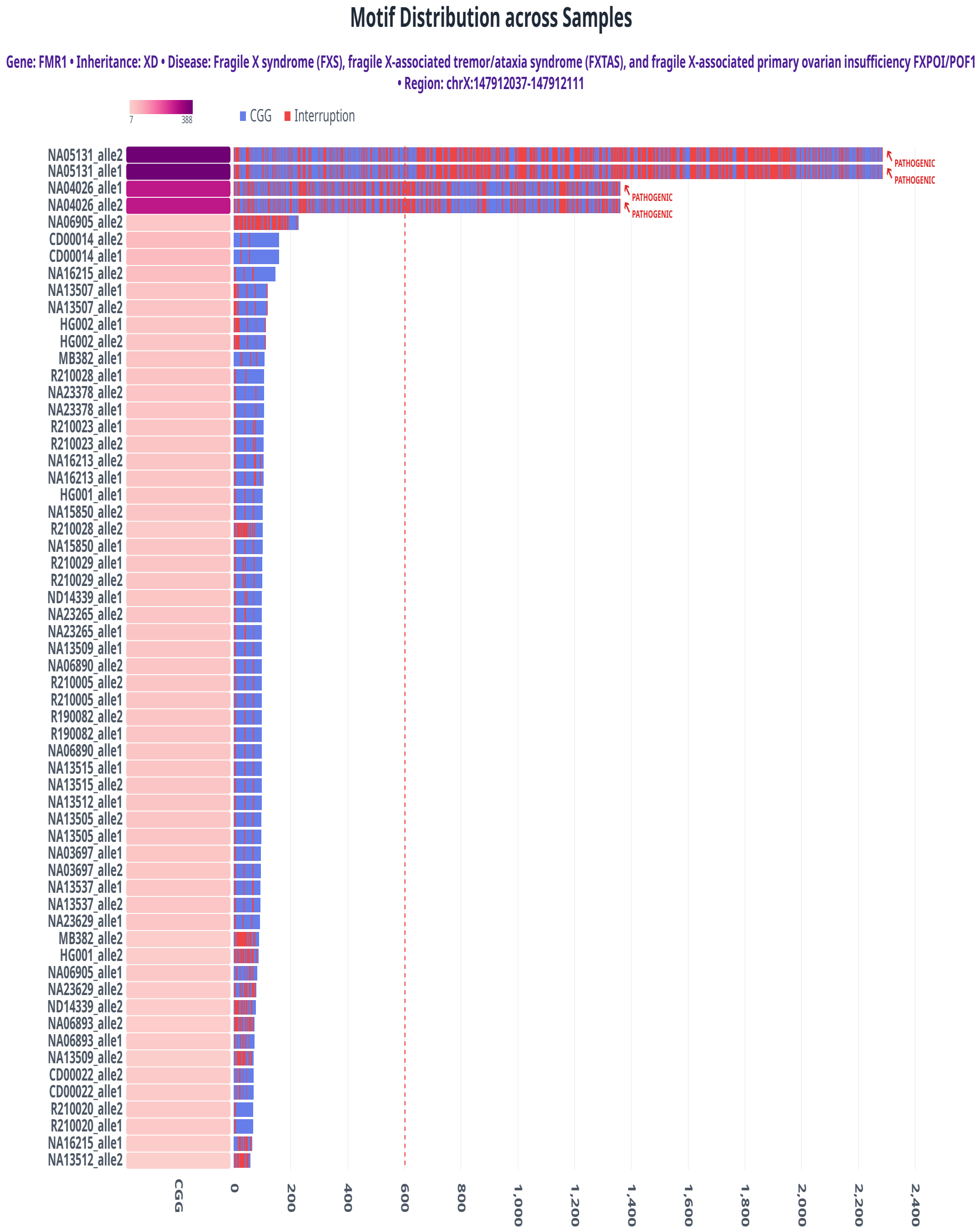


Figure S15: TandemTwister genotyping results for FMR1 gene. Two samples show pathogenic allele expansions. TandemTwister genotyping results for FMR1 gene across 31 samples. Left: Heatmap of motif counts; Right: Bar plot showing copy number alleles per sample. Red dashed line indicates pathogenic cutoff. Two samples (four alleles) show pathogenic allele expansions.
